# CRTC1-MAML2 Establishes a PGC1α-IGF1 Circuit that Confers Vulnerability to PPARγ Inhibition

**DOI:** 10.1101/2020.06.03.129668

**Authors:** Adele M. Musicant, Kshitij Parag-Sharma, Weida Gong, Monideepa Sengupta, Arindam Chatterjee, Erin C. Henry, Yi-Hsuan Tsai, Michele C. Hayward, Siddharth Sheth, Renee Betancourt, Trevor G. Hackman, Ricardo J. Padilla, Joel S. Parker, Jimena Giudice, Colin A. Flaveny, David N. Hayes, Antonio L. Amelio

## Abstract

Mucoepidermoid carcinoma (MEC) is a life-threatening salivary gland cancer that is driven primarily by the transcriptional co-activator fusion CRTC1-MAML2. The mechanisms by which the chimeric CRTC1-MAML2 oncoprotein rewires gene expression programs that promote tumorigenesis remain poorly understood. Here, we show that CRTC1-MAML2 induces transcriptional activation of the non-canonical PGC-1α splice variant PGC-1α4, which regulates PPARγ-dependent IGF-1 expression. This mitogenic transcriptional circuitry is consistent across cell lines and primary tumors. CRTC1-MAML2 positive tumors are dominated by IGF-1 pathway activation and small molecule drug screens reveal that tumor cells harboring the fusion gene are selectively sensitive to IGF-1R inhibition. Furthermore, this dependence on autocrine regulation of IGF-1 transcription renders MEC cells susceptible to PPARγinhibition with inverse agonists. These results yield insights into the aberrant co-regulatory functions of CRTC1-MAML2 and identify a specific vulnerability that can be exploited for precision therapy.

## INTRODUCTION

Transcriptional co-regulators (coactivators and corepressors) regulate gene expression primarily by acting as bridges between DNA-bound transcription factors and the basic transcriptional machinery (Rosenfeld et al., 2006). Chromosomal translocations that create oncogenic fusion genes involving transcriptional co-regulators are predicted to cause profound changes to normal developmental, homeostatic, and/or cellular identity programs in cancer (Lee and Young, 2013; Mitelman et al., 2019; Rabbitts, 1994; Tuna et al., 2019).

Mucoepidermoid carcinoma (MEC) is the most common salivary gland malignancy and patients with advanced recurrent or metastatic tumors often suffer from unresectable, lethal disease marked by a 5-year survival rate of <40% (Bell and Hanna, 2012; El-Naggar et al., 2017; McHugh et al., 2012). Salivary MEC tumors can arise within the major or minor salivary glands and are characterized by significant *intra*-tumoral cellular heterogeneity fueled by cancer stem cells that give rise to multiple cell types including epidermoid, mucus, and intermediate cells (Adams et al., 2015; Seethala et al., 2010; Stewart et al., 1945; Volkmann, 1895). The intermediate cells represent a poorly differentiated, proliferative cell type thought to give rise to the terminally differentiated epidermoid and mucus cell populations. Further, the relative proportion of intermediate cells increases in high-grade tumors and correlates with poor prognosis (Batsakis, 1980). Therefore, transcriptional programs that control cellular identity and support tumor cell functions can directly influence tumor grade and therefore disease progression. Genomic characterization of salivary MEC tumors implicates either a recurrent t(11;19) chromosomal translocation resulting in fusion of two transcriptional coactivators — cAMP Regulated Transcriptional Coactivator 1 (CRTC1) fused to Mastermind-Like 2 (MAML2) — to generate the oncogenic coactivator fusion CRTC1-MAML2 (fusion-positive MEC), or mutations in the tumor suppressor p53 (fusion-negative MEC) (El-Naggar et al., 1996; Kang et al., 2016; Tonon et al., 2003; Wang et al., 2017). The majority of MEC cases are fusion-positive (50-85%) (O’Neill, 2009), and these tumors harbor a strikingly low somatic mutational burden indicating that the CRTC1-MAML2 fusion is the primary oncogenic driver event. Numerous other examples exist of cancers driven primarily by gene fusions in the absence of high mutational burden (Gao et al., 2018; Kadoch and Crabtree, 2013; Missiaglia et al., 2012; Riggi et al., 2014). Although surgical resection is often sufficient to treat patients with low-grade, fusion-positive tumors, some patients expressing CRTC1-MAML2 develop high-grade tumors that display recurrent, chemoradiation-resistant disease (Chen et al., 2007; Seethala and Chiosea, 2016; Warner et al., 2013). This underscores the critical need to develop targeted therapeutic strategies for this subset of salivary MEC patients.

Molecular properties of the chimeric oncoprotein CRTC1-MAML2 have been extensively characterized and reveal that the t(11;19) chromosomal translocation fuses the coiled-coil domain of CRTC1, which promotes binding to the transcription factor cAMP Response Element Binding protein (CREB), with the strong transcriptional activation domain of MAML2 (Coxon et al., 2005; Wu et al., 2005). Consequently, CRTC1-MAML2 functions as a rogue coactivator of the transcription factor CREB, however, its regulatory properties unexpectedly also include gain-of-function interaction and activation of the master transcription factor and proto-oncogene MYC (Amelio et al., 2014). Unfortunately, the absence of ligand binding sites on these transcription factors renders them impractical targets for developing selective inhibitors. Thus, efforts have been directed towards defining the downstream pathways reprogrammed by CRTC1-MAML2. While candidate CREB and MYC target genes have been identified (Amelio et al., 2014; Chen et al., 2015), the subordinate transcriptional pathways dysregulated by aberrant activation of these transcription factors that contribute to transformation of the salivary gland remain poorly defined. Here we performed transcriptomic profiling of fusion positive salivary MEC tumors and report that CRTC1-MAML2 initiates a transcriptional cascade that eventuates with upregulation of the potent growth hormone insulin-like growth factor 1 (IGF-1) via induction of a PGC-1αcoactivator alternative splice variant, PGC-1α4. This has functional consequences for the growth, survival, and oncogenic transformation of salivary gland precursors. Elucidating this pathway establishes a molecular basis for the unique capacity of the CRTC1-MAML2 coactivator fusion to control complex and extensive transcriptional networks beyond CREB and MYC, providing the first evidence of a role for the PGC-1α4 splice variant in human cancer pathogenesis. Integrating drug screening and mechanistic data exposed multiple biologic nodes of drug sensitivity, revealing a CRTC1-MAML2 fusion positive MEC subtype-selective therapeutic vulnerability to PPARγ inhibition.

## RESULTS

### Activation of IGF-1 Signaling is a Hallmark of CRTC1-MAML2 Fusion Positive Salivary Mucoepidermoid Carcinoma

IGF-1 receptor (IGF-1R) overexpression and/or ligand-induced activation of downstream signaling occurs in many cancers and is often required by oncogenes to promote transformation and malignancy (Pollak, 2008). Our previous work investigating the pathobiology of CRTC1-MAML2 positive MEC revealed that several genes involved in anabolic, pro-growth receptor tyrosine kinase (RTK) pathways including IGF-1R signaling are induced by the CRTC1-MAML2 fusion oncogene (Amelio et al., 2014). Here, we confirmed that *IGF-1* is upregulated >100 fold upon ectopic induction of *CRTC1-MAML2* expression (Table S1). To validate the clinical relevance of this finding, we first obtained a cohort of eighteen human primary salivary MEC samples from the surgical pathology department at UNC Hospitals (Chapel Hill, NC) and assessed CRTC1-MAML2 oncogene fusion status (Figure 1A). In accordance with previous reports (Birkeland et al., 2017), we found that ten of these samples (56%) are CRTC1-MAML2 fusion positive. Next, we performed RNA sequencing on all of these MEC samples (fusion positive and negative) to identify differentially expressed genes (DEGs) relative to six normal salivary gland controls (Table S2). Unbiased and unsupervised hierarchical clustering reveal that CRTC1-MAML2 positive tumors constitute a gene expression subtype distinct from CRTC1-MAML2 negative tumors (Figure S1A). Notably, we identified IGF-1 among 3,971 up-regulated genes that are significantly differentially expressed (fold change > 2 and padj < 0.05) in CRTC1-MAML2 fusion positive tumors relative to normal salivary glands (Figure 1B and Table S3). Gene set enrichment analysis (GSEA) of curated gene lists within the Molecular Signatures Database (MSigDB) revealed that IGF signaling pathways rank among the top signatures associated with CRTC1-MAML2 fusion positive tumors (Figure S1B). Thus, we generated a refined gene list (Musicant_MEC_CRTC1-MAML2_IGF1) by curating IGF-1 pathway-related genes with a fold change > 2 (padj<0.05) that reflect a gene signature specific to CRTC1-MAML2 positive MEC tumors (Figures 1C, 1D, S1B, and Table S4). We then evaluated the ability of our curated Musicant_MEC_CRTC1-MAML2_IGF1 gene set to distinguish CRTC1-MAML2 positive MEC by comparison with an established MSigDB GNF2_IGF1 gene set (Figures S1C and S1D). Importantly, we examined expression in these cases by qPCR and found that *CRTC1-MAML2* copy number is significantly correlated (r^2^ = 0.7251; p < 0.0001) with *IGF-1* expression (Figures 1E). Therefore, these results demonstrated that CRTC1-MAML2 fusion positive MEC is characterized by increased IGF-1 expression and suggest an important role for IGF-1R signaling in this MEC tumor subtype.

**Figure 1.**
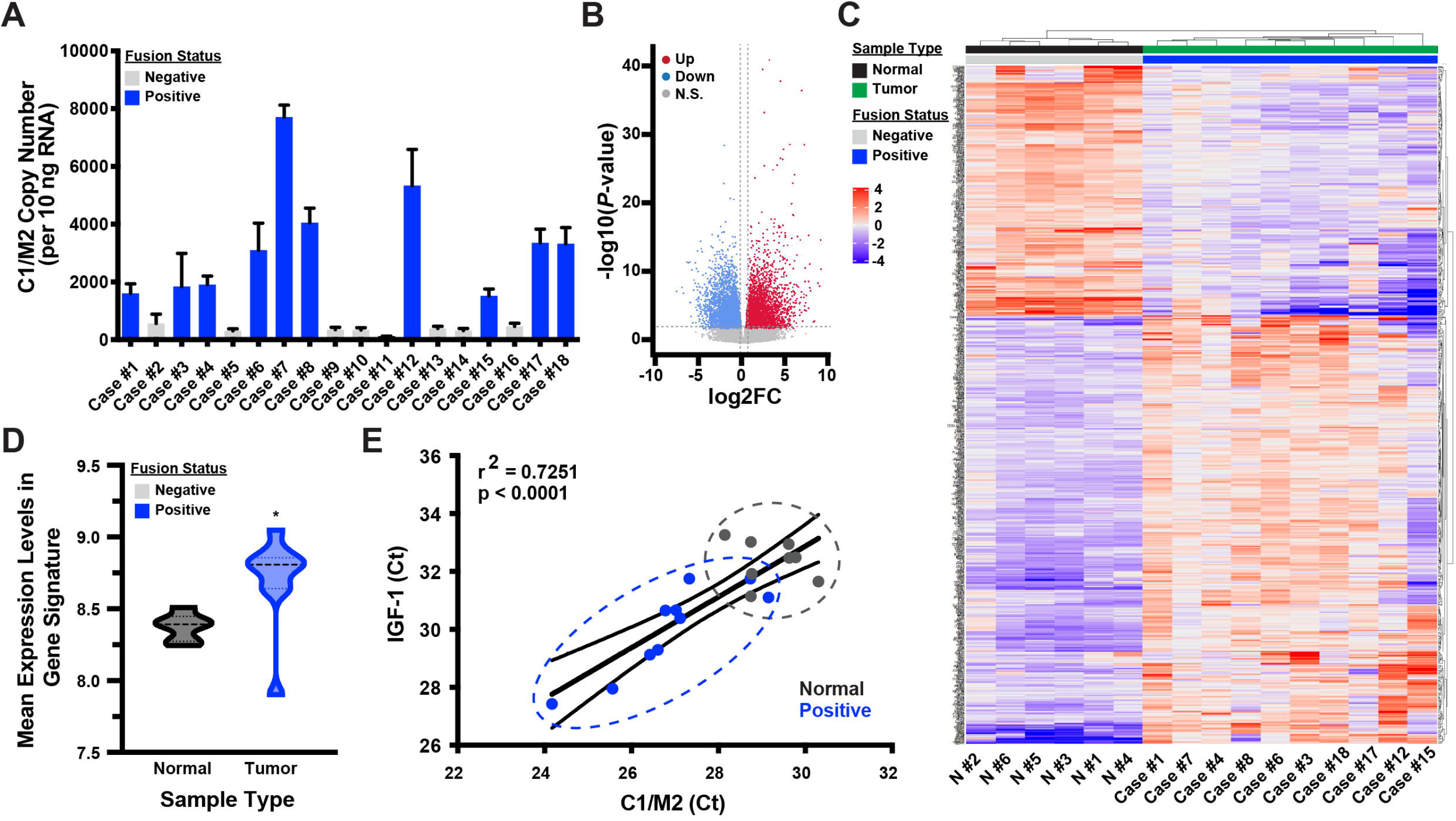
Bioinformatics Analyses Identify IGF-1 as a Hormone Associated with C1/M2 Fusion Positive Salivary MEC. (A) Quantitative real-time PCR analysis of *C1/M2* expression in human salivary MEC tumor samples. *C1/M2* copy number per 10 ng input RNA was calculated based on a standard curve. Samples with <500 *C1/M2* transcripts per 10 ng RNA were classified as fusion-negative (grey bars), while those with ≥500 *C1/M2* transcripts per 10 ng RNA were classified as fusion-positive (blue bars). (B) RNA sequencing was performed on ten C1/M2-positive MEC tumor samples and six normal salivary gland samples. The volcano plot shows genes that are significantly up-regulated (red) and down-regulated (blue) in C1/M2-positive MEC compared to normal salivary glands (padj < 0.05). (C) Heat map of IGF1-related DEGs between fusion-positive MEC and normal salivary gland samples. Normal (Black - e.g. N #1, N #2) and tumor (Green - e.g. Case #1, Case #7) samples are indicated at the top of the heat map in black and green, respectively. C1/M2 fusion status is indicated in grey (Negative) and blue (Positive). (D) Violin plot highlighting the significance of IGF-1 pathway-related genes within our curated Musicant_MEC_CRTC1-MAML2_IGF1 gene set between fusion positive MEC and normal salivary gland samples (*p < 0.05). (E) Comparison of *C1/M2* and *IGF-1* CT values in qPCR data from human MEC tumor (blue) and normal salivary gland (grey) samples. The fitted regression line demonstrates the correlation between the expression levels of the two genes (r^2^ = 0.7251, p < 0.0001). 95% confidence intervals are indicated by curved lines on either side of the linear regression. See also Figure S1.

**Table 1.**
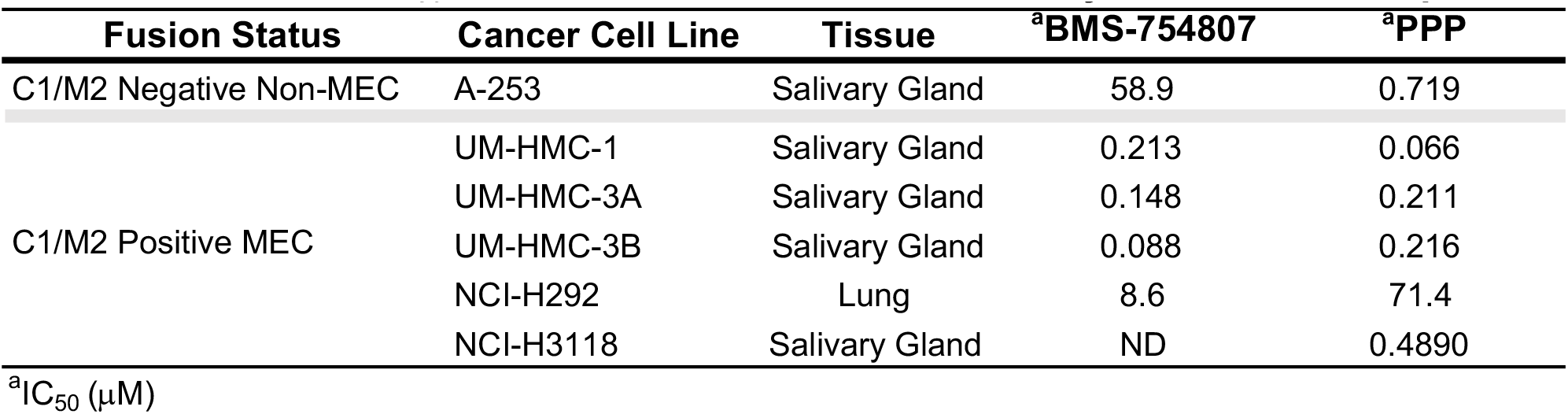
Calculated IC_50_ of IGF-1R inhibitors on the viability of C1/M2 fusion positive and negative cancer cell lines.

**Table 2.** Calculated IC_50_ of PPARγ inverse agonists on the viability of C1/M2 fusion positive and negative cancer cell lines.

| **Cancer Cell Line** | **<sup>a</sup>SR-10221** | **<sup>a</sup>SR-2595** | **<sup>a</sup>T0070907** |
| --- | --- | --- | --- |
| UM-HMC-1 | 114.5 | 40.1 | 328.1 |
| UM-HMC-3A | 72.7 | 50.2 | 222.7 |
| UM-HMC-3B | 148.4 | 78.6 | 329.7 |
| NCI-H292 | 76.6 | ND | ND |
| NCI-H3118 | 98.5 | ND | ND |

<sup>a</sup>IC<sub>50</sub> ( μM)

### CRTC1-MAML2 Fusion Positive MEC Tumor Cell Lines Display Selective Molecular Sensitivities to Targeted IGF-1R Inhibition

To investigate the role of constitutive IGF-1 expression and downstream signaling in CRTC1-MAML2 positive MEC cells, we obtained a panel of five MEC (H292, H3118, HMC1, HMC3A, and HMC3B) and three epidermoid (A253, A388, and A431) cell lines. We first confirmed that all of the MEC cell lines are positive for expression of the *CRTC1-MAML2* fusion transcript, and that all of the epidermoid carcinoma cell lines are fusion negative (Figure 2A). Further, all CRTC1-MAML2 positive MEC cell lines display robust *IGF-1* expression relative to CRTC1-MAML2 negative cell lines, and this increased *IGF-1* expression significantly correlates (r^2^ = 0.8664; p = 0.0008) with *CRTC1-MAML2* expression in fusion positive MEC cells (Figure 2B and S2A). Finally, histologic analysis of tumor xenografts confirm that the CRTC1-MAML2 positive salivary MEC cell lines generate significant *intra*-tumoral cellular heterogeneity, which mimic the mucus, epidermoid, and intermediate cell types characteristic of human salivary MEC (Behboudi et al., 2006; Bell and Hanna, 2012; Warner et al., 2013). In contrast, the fusion negative epidermoid carcinoma cell lines generate xenograft tumors with typical solid epidermoid morphology (Figure 2C). Importantly, immunohistochemical staining demonstrate that CRTC1-MAML2 positive tumors are associated with increased IGF-1 production *in vivo* (Figures 2C and S2B).

**Figure 2.**
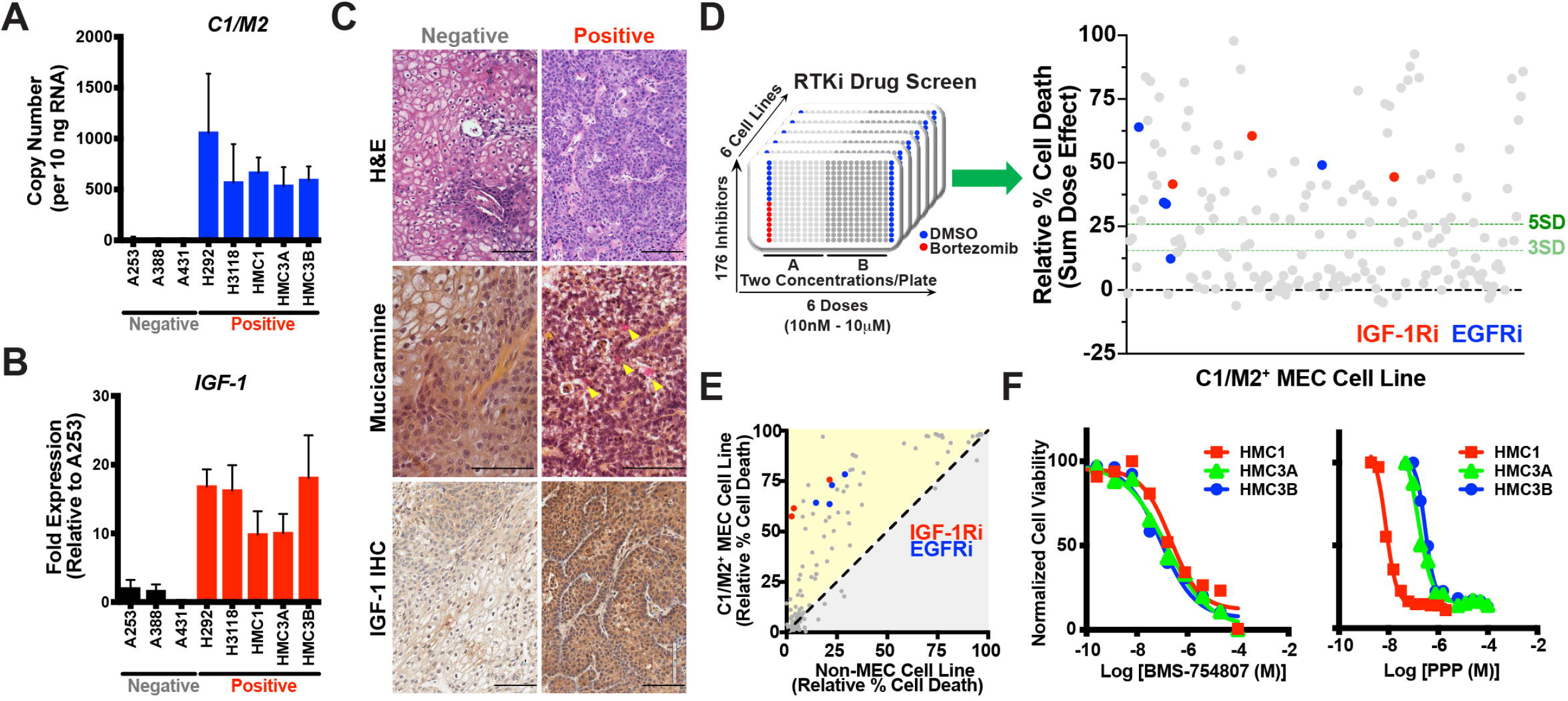
A Focused Small Molecule Drug Screen Identifies Sensitivity of C1/M2 Fusion Positive Tumor Cells to Compounds Targeting IGF-1R. (A) Quantitative real-time PCR analysis of *C1/M2* expression in MEC cell lines and control epidermoid carcinoma cell lines. *C1/M2* copy number per 10 ng input RNA was calculated based on a standard curve. Cell lines are listed along the x-axis, with fusion status (C1/M2-positive or -negative) indicated below in grey (negative) or red (positive). (B) Quantitative real-time PCR analysis of *IGF-1* expression in C1/M2-positive and C1/M2-negative cell lines. Cell lines are listed along the x-axis, with fusion status (C1/M2-positive or -negative) indicated below in grey (negative) or red (positive). Relative fold expression is shown normalized to *RPL23* mRNA levels. (C) Histologic and immunohistochemical analysis of C1/M2-positive and C1/M2-negative xenografts. Representative xenograft sections were formalin fixed, embedded, and H&E stained (top). Mucicarmine staining (middle) revealed mucus cells (pink staining, indicated by yellow arrowheads) only in C1/M2-positive xenografts. IHC staining for IGF-1 (bottom) demonstrated that IGF-1 expression is increased in C1/M2-positive xenografts relative to C1/M2-negative xenografts. Scale bar, 100 μm. (D) A drug screen was performed in six cell lines (five C1/M2-positive MEC cell lines and one C1/M2-negative epidermoid carcinoma cell line). 176 inhibitors were tested, in duplicate, across all cell lines at six concentrations ranging from 10 nM to 10 µM. DMSO (1%) and Bortezomib (1 µM) were used as negative and positive controls, respectively, on each plate. Several IGF-1R and EGFR inhibitors emerged as top hits, inducing cell death in C1/M2-positive cell lines at >5 standard deviations above baseline (DMSO). (E) Relative cell death induced by drug screen inhibitors. IGF1R and EGFR inhibitors more effectively induced cell death in C1/M2-positive MEC cells compared with C1/M2-negative epidermoid carcinoma cells. (F) Representative dose response curves showing viability of three C1/M2-positive cell lines (HMC1, HMC3A, and HMC3B) treated with increasing concentrations of IGF1R inhibitors (BMS-754807 and PPP). *See also Table 1 for a summary of the IC50 values*. See also Figures S2-S3.

To test whether CRTC1-MAML2 positive MEC cells are dependent on IGF-1 signaling, we performed an unbiased small molecule drug screen focused on receptor tyrosine kinase inhibitors (RTKi) at concentrations ranging from 10 nM to 10 µM in all five CRTC1-MAML2 fusion positive cells lines relative to a fusion negative cell line control (Figure 2D, *left*). Calculation of the Z’-factor demonstrated robust assay quality such that the distribution between the positive (Bortezimib) and negative (DMSO) controls indicates low likelihood of false positive hits (Z’ > 0.75) (Figure S3A). Multiple independent inhibitors targeting IGF-1R and EGFR emerged as significant hits (p ≥ 3SD) in this screen (Figure 2D, *right*). Notably, EGFR has been explored as a therapeutic target although with limited clinical success (Chen et al., 2015). Compared to the non-MEC fusion negative control cells, IGF-1R inhibitors induced selective and robust cell death in all CRTC1-MAML2 positive cell lines (Figure 2E, S3B, and S3C), indicating the relative importance of IGF-1/IGF-1R signaling in CRTC1-MAML2 fusion positive MEC. Dose titrations of two separate IGF-1R inhibitors, BMS-754807 and picropodophyllin (PPP), confirmed that the CRTC1-MAML2 cell lines are sensitive to IGF-1R inhibition with IC50s for both compounds in the low nM range (Figure 2F and Table 1).

### IGF-1R Signaling is Critical for CRTC1-MAML2 Fusion Positive MEC Tumor Cell Growth and Survival

We next wanted to examine the functional role of IGF-1R signaling on human CRTC1-MAML2 fusion positive MEC cell growth and survival. Live-cell, kinetic proliferation assays revealed that wild-type HMC3A cells reach 50% confluency after 28 hr (GC50 = 28 hr), but pharmacologic inhibition of IGF-1R with PPP increases this time to confluency ∼36% relative to control (Figure 3A). Similarly, genetic inhibition of IGF-1R by shRNA-mediated knockdown (validated in Figures S4A and S4B) also significantly increased the GC50 ∼74% to 48 hr relative to control. Remarkably, the combination of pharmacologic and genetic inhibition of IGF-1R dramatically increased the GC50 ∼94% to 54 hr (Figure 3A). To test if blocking IGF-1R impairs the tumorigenic potential of MEC cells to grow as clonogenic colonies *ex vivo*, CRTC1-MAML2 positive cells were plated at low density and observed for colony formation. We found that both PPP and BMS-754807 markedly blunted the clonogenic potential of several independent MEC cell lines relative to vehicle-treated cells (Figures 3B and S4C). Several recent studies have documented the existence of cancer stem cell (CSC) subpopulations within various head and neck cancers including salivary MEC and these CSCs are associated with malignant potential (Adams et al., 2015; Curtarelli et al., 2018; Keysar et al., 2017). Since tumor spheroids are uniquely enriched for CSCs, we also quantified the efficacy of PPP on 3D tumor spheroid formation in Matrigel and find that sustained IGF-1Ri treatment (7 days) significantly blocks MEC 3D sphere formation (Figures 3C and 3D). We also investigated the mechanism underlying the observed decrease in proliferation and tumorigenic potential and found that PPP stimulates a significant dose-dependent increase (p < 0.0001) in apoptosis compared to vehicle-treated cells (Figure 3E).

**Figure 3.**
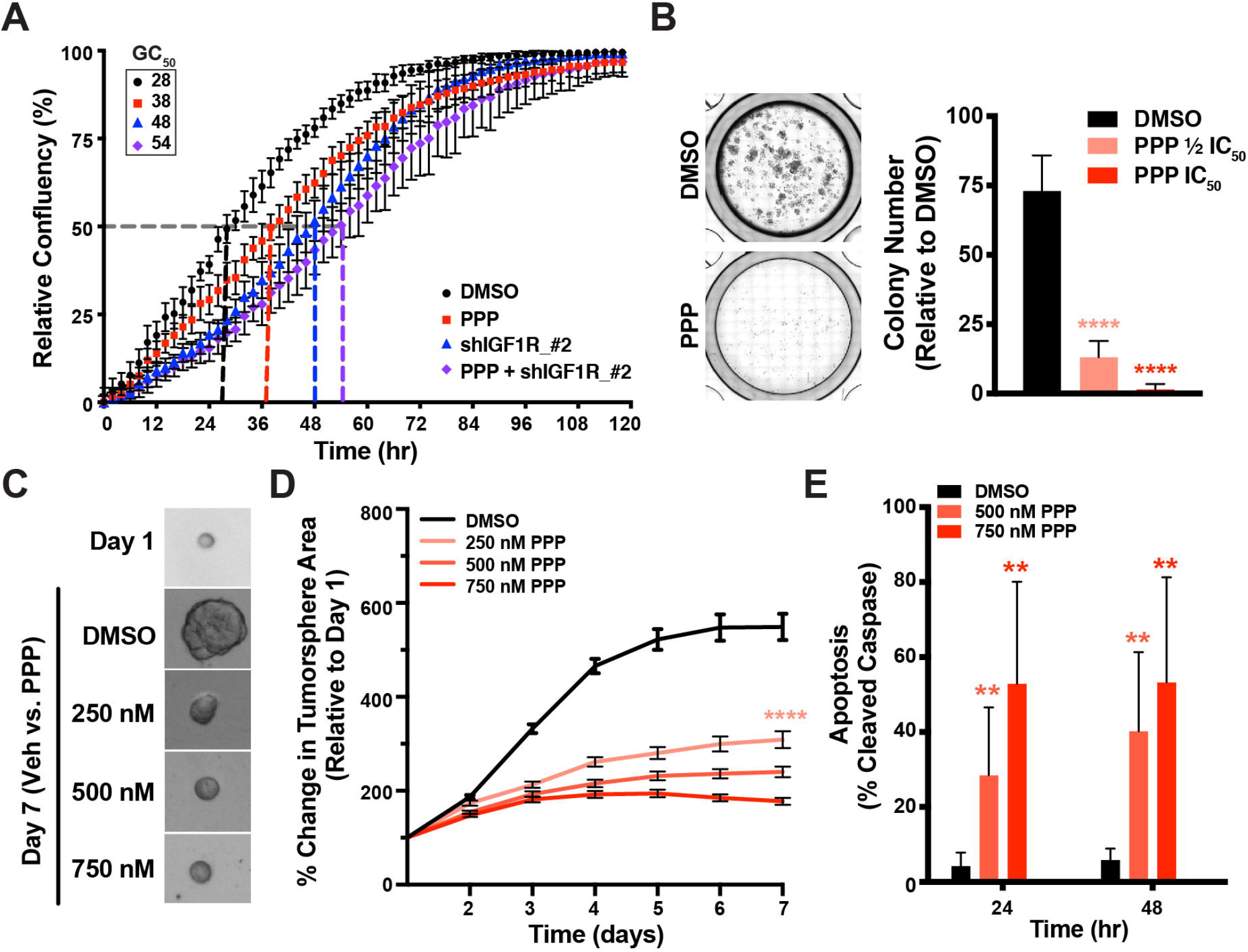
Inhibition of IGF-1R Blocks MEC Cell Growth and Induces Apoptosis. (A) Cell proliferation assay showing relative confluency of HMC3A cells treated with DMSO or PPP (IGF-1R inhibitor) with and without shRNA-mediated *IGF-1R* knockdown. GC50 values for each condition are shown in hours and indicate the time required for cells to reach 50% confluency. (n = 3 biological replicates, mean ± SEM). (B) Number of colonies formed by HMC3A cells treated with DMSO or PPP at ½ IC50 concentration or IC50 concentration. *See also Table 1 for a summary of the IC50 values*. A colony is defined as a cluster of ≥50 cells. Representative images of wells treated with DMSO and PPP (IC50 concentration) are shown to the left (n = 3 biological replicates, mean ± SEM). \*\*\*\**p* ≤ 0.0001. (C) Representative images of HMC3A tumorspheres on day one (top) and seven days post treatment with DMSO (vehicle control) or increasing concentrations of PPP. (D) Tumorsphere formation in HMC3A cells treated with DMSO or increasing concentrations of PPP. Percent change in tumorsphere area is calculated as [TumorAreaDayX]/[TumorAreaDay1]*100. >50 individual tumorspheres per condition were tracked for each individual experiment. (n = 3 biological replicates, mean ± SEM). \*\*\*\**p* ≤ 0.0001. (E) Apoptosis levels (measured as % cleaved Caspase 3/7) in HMC3A cells treated with DMSO or PPP. Data was collected via flow cytometry and normalized to a non-stained control for each condition. (n = 3 biological replicates, mean ± SEM). \*\**p* ≤ 0.01. See also Figure S4.

### CRTC1-MAML2 Establishes a Synthetic PGC-1α4 Circuit that Regulates IGF-1 Expression in MEC Tumor Cells

To determine the mechanism by which CRTC1-MAML2 regulates *IGF-1* expression in MEC cells, we utilized our engineered CRTC1-MAML2-inducible stable cell line (doxycycline-regulated HEK293-*CMV^TetR^TetO^C1/M2^*) (Amelio et al., 2014) (Figure S5A). Overexpression of CRTC1-MAML2 in this cell line significantly increases *IGF-1* expression at both the transcript and protein levels (Figures 4A, 4B, S5B, and S5C). CRTC1-MAML2 coactivates the transcription factor CREB, which in turn binds to CREB at specific DNA sequences called cAMP-response elements (CREs). Examination of IGF-1 promoter sequences revealed a non-canonical CRE site, indicating that CRTC1-MAML2 could possibly drive IGF-1 signaling through direct upregulation of *IGF-1* transcription. However, chromatin immunoprecipitation (ChIP) assays failed to identify CRTC1-MAML2 enrichment at the *IGF-1* promoter relative to the control, canonical CREB target gene *NR4A2* (Figure 4C).

**Figure 4.**
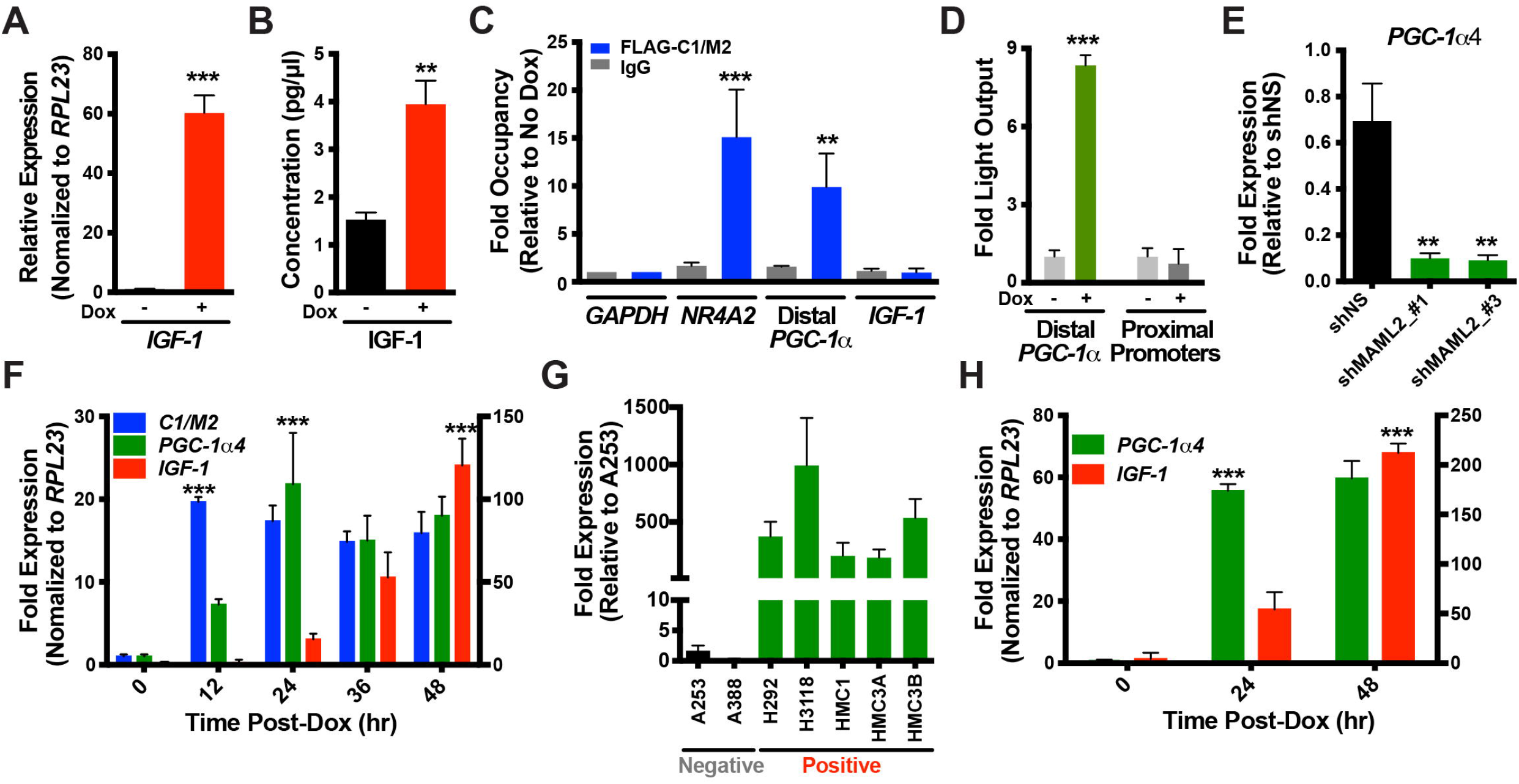
C1/M2 Fusion Induces Expression of the PGC-1α4 Alternative Splice Variant Leading to IGF-1 Upregulation. (A) Quantitative real-time PCR analysis of *IGF-1* mRNA levels in C1/M2-inducible cells (doxycycline-regulated HEK293-*CMV^TetR^TetO^C1/M2^* cells) treated with and without 1 µg/mL doxycycline. Relative fold expression is shown normalized to *RPL23* mRNA levels. (n = 3 biological replicates, mean ± SEM). ***p < 0.001. (B) AlphaLISA-based analysis of secreted IGF-1 in media from C1/M2-inducible cells (HEK293-*CMV^TetR^TetO^C1/M2^*) with and without 1 µg/mL doxycycline treatment. (n = 2 biological replicates, mean ± SEM). **p = 0.003. (C) Chromatin immunoprecipitation analysis of Flag-C1/M2 promoter occupancy in C1/M2-inducible cells (HEK293-*CMV^TetR^TetO^C1/M2^*). Data are expressed as promoter occupancy in doxycycline-treated cells normalized to promoter occupancy in vehicle-treated cells. GAPDH and NR4A2 promoters were used as negative and positive controls, respectively. (n = 3 biological replicates, mean ± SEM). ***p < 0.001, **p < 0.01. (D) Luciferase assay measuring activation of the distal and proximal PGC-1α promoters in C1/M2-inducible cells (HEK293-*CMV^TetR^TetO^C1/M2^*) with and without 1 µg/mL doxycycline treatment. (n = 3 biological replicates, mean ± SEM). ***p < 0.001. (E) Quantitative real-time PCR analysis of *PGC-1α4* mRNA expression in HMC3A cells with and without shRNA-mediated *C1/M2* knockdown. Relative fold expression is shown normalized to *RPL23* mRNA levels and to mock transduced condition. shNS: non-specific shRNA. shMAML2_#1 and shMAML2_#3: shRNAs targeting *C1/M2* and *MAML2*. (n = 2 biological replicates, mean ± SEM). **p < 0.01. (F) Temporal analysis of *C1/M2*, *PGC-1α4*, and *IGF-1* expression kinetics in C1/M2-inducible cells (HEK293-*CMV^TetR^TetO^C1/M2^*) treated with 1 µg/mL doxycycline for 0-48 hours. *C1/M2* and *PGC-1α4* expression is indicated on the left y-axis, while *IGF-1* expression is indicated on the right y-axis. Relative fold expression is shown normalized to *RPL23* mRNA levels. (n = 3 biological replicates, mean ± SEM). ***p < 0.001. (G) Quantitative real-time PCR analysis of *PGC-1α4* expression in C1/M2-positive and C1/M2-negative cell lines. Cell lines are listed along the x-axis, with fusion status (C1/M2-positive or -negative) indicated below in grey (negative) or red (positive). Relative fold expression is shown normalized to *RPL23* mRNA levels. (n = 3 biological replicates, mean ± SEM). (H) Quantitative real-time PCR analysis of *PGC-1α4* and *IGF-1* mRNA expression in *PGC-1α4*-inducible cells (doxycycline-regulated HEK293-*PGK^TetOn3G^TRE3GS^PGC-1α4^* cells) treated with 1 µg/mL doxycycline for 0-48 hours. Relative fold expression is shown normalized to *RPL23* mRNA levels. (n = 3 biological replicates, mean ± SEM). ***p < 0.001. See also Figure S5.

We previously showed that CREB regulates stress-inducible expression of a PPARγ coactivator 1αsplice variant (PGC-1α4) transcribed from an upstream (∼13 Kbp) distal promoter of the PGC-1α locus (Bruno et al., 2014), which in turn was shown to selectively regulate *IGF-1* expression (Ruas et al., 2012). Thus, we performed ChIP assays and identified CRTC1-MAML2 enrichment specifically at this distal PGC-1α promoter (Figure 4C). Moreover, luciferase reporter assays confirmed that CRTC1-MAML2 activates the distal PGC-1α promoter but not the proximal promoter (Figure 4D), suggesting that CRTC1-MAML2 may indirectly regulate *IGF-1* expression by controlling *PGC-1α4* expression in MEC cells. We next performed shRNA-mediated *CRTC1-MAML2* knockdowns and found that *PGC-1α4* levels are dramatically reduced similar to the control *NR4A2*, confirming that CRTC1-MAML2 directly upregulates *PGC-1α4* transcription (Figure 4E and S5D). Expression of inducible dominant-negative CREB (A-CREB) in a stable MEC cell line (HMC3A-*PGK^TetOn3G^TRE3GS^A-CREB^*) also blocked *PGC-1α4* expression, supporting the role of CREB-dependent regulation of the *PGC-1α* distal promoter by CRTC1-MAML2 (Figure S5E). To examine the kinetics of this expression circuit more closely, we performed a time course experiment in CRTC1-MAML2-inducible stable cells and found that *PGC-1α4* transcripts peak at 24 hrs (∼12 hr after *CRTC1-MAML2* induction) while *IGF-1* transcripts begin to rise at ∼24-36 hrs, but peak 48 hrs after *CRTC1-MAML2* induction and 24 hrs after *PGC-1α4* upregulation (Figure 4F). Notably, CRTC1-MAML2 positive MEC cell lines are all characterized by robust *PGC-1α4* expression relative to CRTC1-MAML2 negative cell lines (Figure 4G). We also generated a *PGC-1α4*-inducible stable cell line (doxycycline-regulated HEK293-*PGK^TetOn3G^TRE3GS^PGC-1α4^*; validated in Figures S5F and S5G) and confirmed that overexpression of the PGC-1α4 splice variant acts as the intermediate to CRTC1-MAML2, which directly regulates *IGF-1* expression (Figure 4H). Finally, shRNA-mediated knockdown of PGC-1α4, as well as other components within this signaling circuit (IGF-1R and CRTC1-MAML2) significantly blunts proliferation of CRTC1-MAML2 positive MEC cells (Figure S5H), confirming that PGC-1α4 is both necessary and sufficient for activating the IGF-1 signaling circuit in fusion positive MEC.

### The PGC-1α4 Co-Regulator Activates PPARγ-dependent Transcription of IGF-1 in CRTC1-MAML2 Fusion Positive MEC Tumor Cells

PGC-1α is a co-activator that binds to the transcription factor peroxisome proliferator-activated receptor gamma (PPARγ) to exert its effects on transcription of metabolic target genes (Semple et al., 2006; Sonoda et al., 2007). Strikingly, analysis of the *IGF-1* locus identified three canonical PPAR-response element (PPRE) binding motifs within the *IGF-1* promoter region (Figure S6A). To test whether PGC-1α4 co-activates PPARγ to modulate *IGF-1* transcription, we first confirmed that CRTC1-MAML2 induces transcription of an *IGF-1* promoter-driven luciferase reporter comparable to levels of induction observed with a PPRE-driven luciferase reporter upon CRTC1-MAML2 overexpression (Figures S6B and S6C). Importantly, functional characterization revealed that this luciferase reporter construct is responsive to the PPARγagonist GW1929 indicating that the cloned *IGF-1* promoter fragment includes the PPRE motifs that recruit PPARγ (Figure S6D). Therefore, we next tested whether the PPARγ inverse agonist SR10221, which functions to recruit transcriptional co-repressors to PPARγ that repress target gene transcription, are effective at inhibiting *IGF-1* expression. Treatment with SR10221 effectively downregulated PPARγ-mediated transcriptional activity of the *IGF1*-Luciferase reporter in a dose-dependent manner to below basal levels, comparable to that observed with the *PPRE*-Luciferase reporter (Figure 5A). This indicates that the PGC-1α4:PPARγcircuit can control *IGF-1* expression and suggests that drugs targeting PPARγ may provide therapeutic benefit against CRTC1-MAML2 positive salivary MEC.

**Figure 5.**
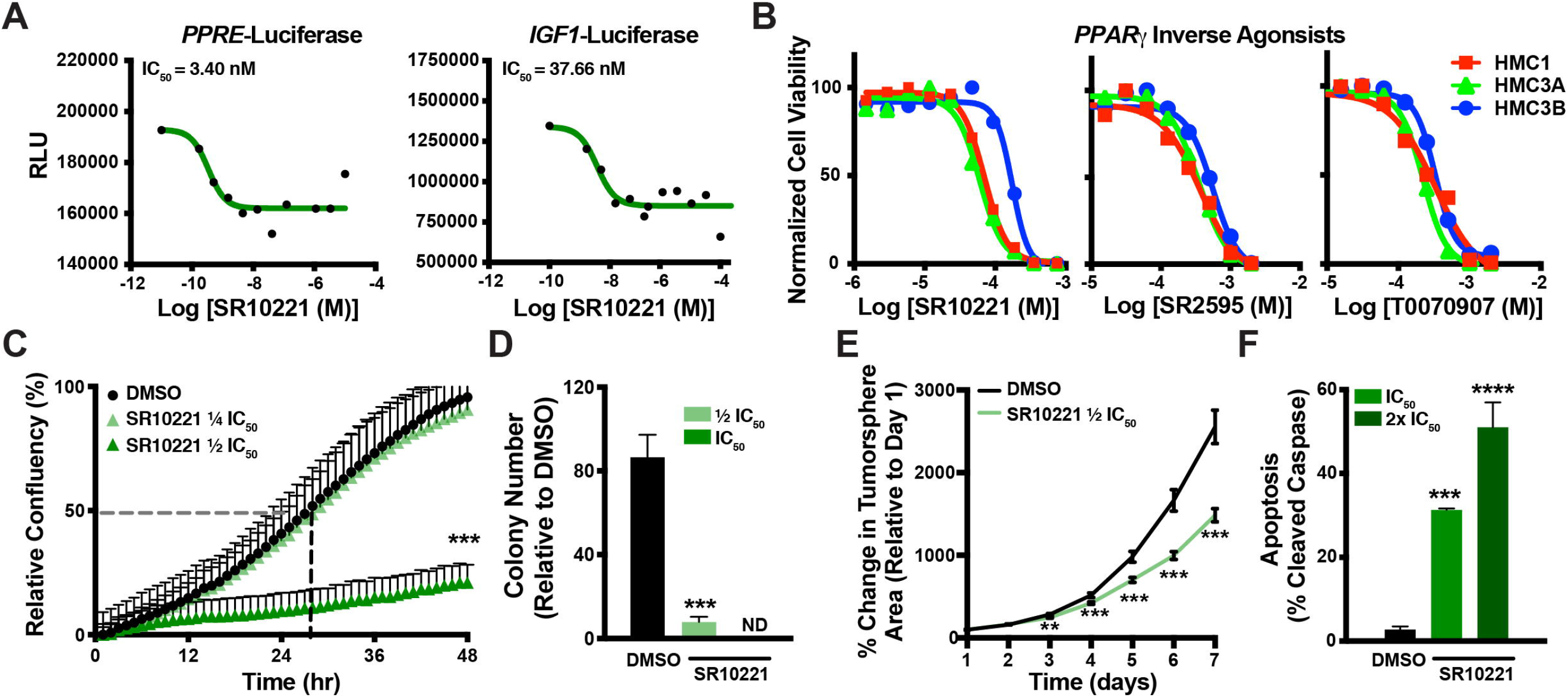
Inhibition of IGF-1 Expression with PPARγ Inverse Agonists Blocks MEC Cell Growth and Induces Apoptosis. (A) PPARγ-response element (PPRE)-driven (*left*) and IGF-1 promoter-driven (*right*) luciferase reporter assay showing repression of basal transcriptional activity of endogenously expressed PPARγ in HEK293-*CMV^TetR^TetO^C1/M2^* cells expressing C1/M2. Cells were treated with increasing concentrations of the PPARγ inverse agonist SR10221 for 24 hr prior to measuring luciferase activity (n = 2 biological replicates, mean ± SEM). (B) Representative dose response curves showing viability of three C1/M2-positive cell lines (HMC1, HMC3A, and HMC3B) treated with increasing concentrations of PPARγ inverse agonists (SR10221, SR2595, and T0070907). (n = 2-3 biological replicates, mean ± SEM). *See also Table 2 for a summary of the IC50 values*. (C) Cell proliferation assay showing relative confluency of HMC3A cells treated with DMSO or SR10221 at ½ IC50 concentration or ¼ IC50 concentration (*see Table 2 for IC50 values*). (n = 3 biological replicates, mean ± SEM). (D) Colony formation in HMC1 cells treated with DMSO or SR10221 at ½ IC50 concentration or IC50 concentration (*see Table 2 for IC50 values*). A colony is defined as a cluster of ≥50 cells (n = 3 biological replicates, mean ± SEM). \*\*\**p* ≤ 0.001. (E) Tumorsphere formation in HMC3A cells treated with DMSO or ½ IC50 SR10221. Percent change in tumorsphere area is calculated as [TumorAreaDayX]/[TumorAreaDay1]*100. >50 individual tumorspheres per condition were tracked for each individual experiment. (n = 3 biological replicates, mean ± SEM). \*\**p* ≤ 0.01, \*\*\**p* ≤ 0.001. (F) Apoptosis levels (measured as % cleaved Caspase 3/7) in HMC3A cells treated with DMSO or SR10221 at IC50 concentration or 2x IC50 concentration (*see Table 2 for IC50 values*). Data was collected via flow cytometry and normalized to a non-stained control for each condition. (n = 3 biological replicates, mean ± SEM). \*\*\**p* ≤ 0.001, \*\*\*\**p* ≤ 0.0001. See also Figure S6.

### PPARγ Inverse Agonists Inhibit IGF-1 Expression and Suppress CRTC1-MAML2 Fusion Positive MEC Tumor Growth

To investigate the functional role of PPARγ dependency on MEC cell growth and survival, we tested the effects of several PPARγ inverse agonists in our panel of CRTC1-MAML2 positive MEC cell lines. First, to determine the efficacy of PPARγ inverse agonists as anti-MEC agents, we assessed the effect of three different inverse agonist (SR10221, SR2595, and T0070907) on cancer cell viability using the full panel of CRTC1-MAML2 positive MEC cell lines (Figure 5B and Table 2). Dose titration of these compounds revealed that SR10221 potently reduced cancer cell viability at low micromolar concentrations (half-maximal inhibitory concentration [IC_50_] ∼70 – 150 μM) in HMC3A cells (Table 2). Treatment of CRTC1-MAML2 positive cell lines with sub-lethal doses of SR10221 (½ IC50) significantly decreased proliferation, 2D colony formation, and 3D tumor spheroid growth (Figures 5C, 5D, 5E, and S6E). Moreover, this decreased growth and tumorigenic potential coincided with potent induction of apoptotic cell death as evidenced by significant increases in caspase-3/7 activation (Figure 5F).

To determine the efficacy of PPARγ inverse agonists *in vivo*, we first tested the potency of SR10221 in subcutaneous HMC3A tumor xenograft models labeled with our LumiFluor bioluminescent reporter (Schaub et al., 2015). Tumors xenografts were allowed to grow to a palpable size (approximately 50 mm^3^) and animals were randomized into two cohorts. These cohorts then received *intra*-peritoneal (*i.p.*) administration of 20 mg/kg SR10221 or vehicle control once daily for three weeks which resulted in the significant inhibition (p < 0.0001) of HMC3A tumor xenograft growth (Figure 6A and 6B). Similarly, we also tested the *in vivo* potency of SR2595 since it displays superior pharmacokinetic properties, although we administered SR2595 at 60 mg/kg since SR10221 was shown to possess ∼2- to 3-fold greater potency *in vitro* (Marciano et al., 2015). Treatment with SR2595 at this concentration was generally well-tolerated without body weight loss (Figure 6D), but significantly blocked the growth of CRTC1-MAML2 positive MEC tumor xenografts (Figure 6C, 6E, 6F, and 6G). Collectively, these results suggest that inhibition of IGF-1 signaling using PPARγ inverse agonists alone or in combination with IGF-1R inhibitors may be a viable therapeutic strategy for the targeted treatment of CRTC1-MAML2 positive MEC.

**Figure 6.**
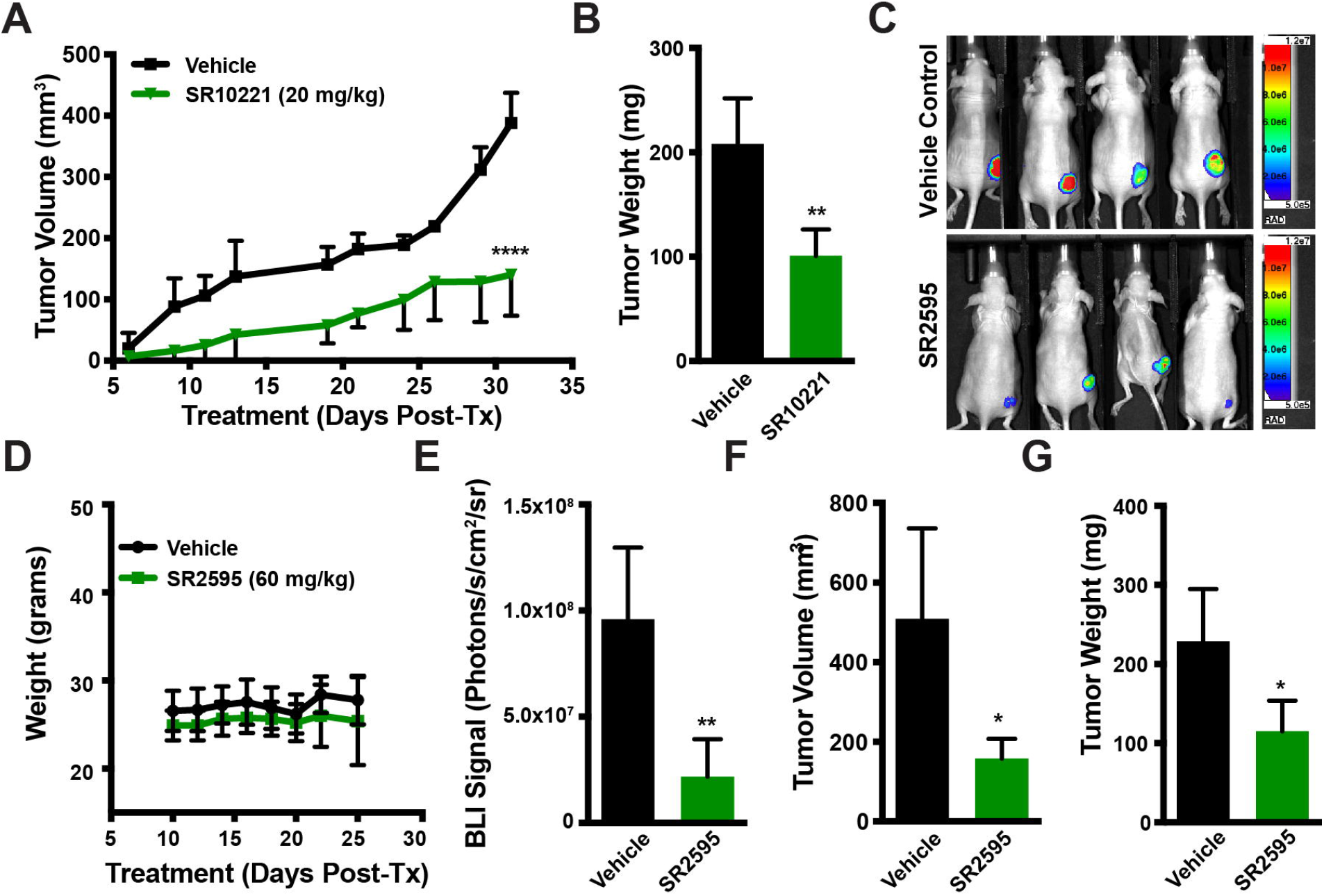
PPARγ Inverse Agonists Suppress Tumor Growth in Xenograft Models of Salivary MEC. (A) Growth of HMC3A xenograft tumors over time during treatment with either SR10221 or a vehicle control. Tumor xenograft volume was quantified by caliper measurements obtained on the indicated days following initiation of drug administration. Decreased tumor volume in the SR10221 cohort correlates directly with suppression of tumor growth *in vivo* (n = 4, mean ± SEM). \*\*\*\**p* ≤ 0.0001. (B) Weight of resected vehicle- and SR10221-treated HMC3A tumor xenografts at study endpoint. \*\**p* ≤ 0.01. (C) Representative *in vivo* bioluminescent images obtained at endpoint of Nu/Nu mice bearing subcutaneous HMC3A tumor xenografts and treated with either SR2595 or a vehicle control. Images were captured with an open filter and an acquisition time of 5 min (binning = 2; FOV = 25; Fstop = 2; object height = 1.5). (D) Weights of Nu/Nu mice bearing subcutaneous HMC3A xenograft tumors and treated with SR2595 or a vehicle control for up to 24 days. (E) Comparison of HMC3A xenograft tumor growth following treatment with either SR2595 or vehicle control. Bioluminescence imaging (BLI) signals from tumor xenografts were quantified by region of interest (ROI) analysis of images obtained at endpoint (day 24 following initiation of drug administration). Decreased BLI signal in the SR2595 cohort correlates directly with suppression of tumor growth *in vivo* (n = 4, mean ± SEM). \*\**p* ≤ 0.01. (F) Volume of resected vehicle- and SR2595-treated HMC3A xenograft tumors at study endpoint. *p ≤ 0.05. (G) Weight of resected vehicle- and SR2595-treated HMC3A xenograft tumors at study endpoint. *p ≤ 0.05.

## DISCUSSION

Recurrent chromosomal translocations that generate novel gene fusions have long been known to have the potential to function as cancer drivers (Rabbitts, 1994; Rabbitts, 2009; Rowley, 2001). However, the advent of fusion detection algorithms applied to “omics”-level data has not only enabled the discovery of additional gene fusions composed of splicing factors, signal transduction proteins, transcription factors, and/or transcriptional co-regulators (Fernandez-Cuesta et al., 2015; Jang et al., 2020; Kumar-Sinha et al., 2015), but has also aided in elucidating the direct functional consequences of these fusions on cellular processes such as signaling and gene expression (Latysheva et al., 2016; Lee and Young, 2013). Interestingly, these aberrant processes are frequently unique to distinct gene fusions which may in turn be characteristic of specific cancer subtypes (Fishbein et al., 2017; Gao et al., 2018). CRTC1-MAML2 represents one such gene fusion composed of two transcriptional co-activators that is pathognomonic for mucoepidermoid carcinoma (MEC) (O’Neill, 2009; Tonon et al., 2003). Here, we report that the CRTC1-MAML2 fusion oncogene directs profound reprogramming of transcriptional networks and establishes a synthetic IGF-1 signal circuit through aberrant expression of an alternative PGC-1α splice variant in salivary MEC.

Despite being the most common salivary gland malignancy, MECs are relatively rare compared to other head and neck cancers and consequently the pathobiology of CRTC1-MAML2 remains poorly understood. While it is clear that patients with advanced stage and/or high-grade tumors display significantly worse overall survival (Chen et al., 2013), the prognostic value of fusion status has recently been challenged (Seethala and Chiosea, 2016). Most MEC patients are treated with surgical excision, which is sometimes accompanied by adjuvant radiotherapy (Nance et al., 2008). Unfortunately, resistance to chemoradiotherapies and a lack of targeted therapies for recurrent/metastatic disease poses significant challenges to treating patients with aggressive MEC tumors. For example, recent studies have indicated that paclitaxel, trastuzumab, or receptor tyrosine kinase inhibitors may be viable treatments for fusion-positive MEC, however preliminary results show only modest responses to these treatments (Coca-Pelaz et al., 2015; McHugh et al., 2012) emphasizing the need for performing thorough molecular profiling studies to aid in the development of new targeted therapies.

As a transcriptional coactivator gene fusion, CRTC1-MAML2 interacts with and activates two master transcription factors, CREB and MYC (Amelio et al., 2014; Tasoulas et al., 2019). Unfortunately, intrinsic disorder and the lack of available binding pockets has made designing small molecules that can effectively target transcription factors a major challenge (Bishop et al., 2019; Chen and Koehler, 2020; Wachtel and Schäfer, 2018). In addition, two recent genomics studies revealed that CRTC1-MAML2 is the primary oncogenic driver in most salivary MECs and that these fusion positive cases lack other cooperating mutations that occasionally serve as actionable targets in other cancers (Kang et al., 2016; Wang et al., 2017). In this study, we set out to identify the most significantly altered and clinically-relevant target genes and pathways downstream of CRTC1-MAML2 that can be exploited to develop novel therapeutic approaches for treating patients with salivary MEC. Using transcriptomic profiling, we show that CRTC1-MAML2 expression is correlated with increased IGF-1 expression and pathway activation in salivary MEC tumors. Autocrine and paracrine activation of the IGF-1 pathway is known to promote growth and survival of multiple tumor types (Pollak, 2008; Yu and Rohan, 2000). Notably, endocrine IGF-1 signaling is broadly involved in normal tissue growth and development through its stimulation of the IGF-1 receptor (IGF-1R) and has been shown to play critical roles in regulating salivary gland growth and development, and in stimulating regeneration following injury (Amano and Iseki, 1993; Kerr et al., 1995; Ryan et al., 1992; Werner and Katz, 2004) thus supporting an important role for aberrant IGF-1 expression in salivary tumor development. Through the use of bioinformatics analyses and unbiased small molecule drug screening, we were successful in pinpointing IGF-1R as an actionable target in salivary MEC suggesting that treatments directed against IGF-1R may provide therapeutic benefit. However, despite early excitement surrounding the potential utility of these treatments for many cancers (Gualberto and Pollak, 2009), targeting the IGF axis has yielded disappointing results in clinical trials (Chen and Sharon, 2013; Denduluri et al., 2015). Thus, the potential utility of anti-IGF-1R antibodies and small molecule drug inhibitors, alone or in combination with other therapeutic modalities is being investigated as a viable alternative approach. Therefore, we also sought to explore the underlying mechanism governing IGF-1 activation by CRTC1-MAML2 in salivary MEC and identified unexpected rewiring of this growth factor signal circuit coordinated by aberrant expression of an alternatively spliced target gene of CRTC1-MAML2/CREB, the PGC-1α4 variant.

PGC-1α has been shown to exert oncogenic (Frattini et al., 2018) or tumor suppressive (Torrano et al., 2016) activity depending on cell type and context-dependent metabolic cues (Mastropasqua et al., 2018; Sancho et al., 2015; Xing et al., 2017). Notably, the PGC-1α family of transcriptional coactivators is composed of an expanding list of transcript isoforms generated by multiple promoters and alternative pre-mRNA splicing, and these resulting protein variants are known to display unique functional properties (Martínez-Redondo et al., 2016; Martínez-Redondo et al., 2015). Rare, alternatively spliced transcripts such as PGC-1α4 are emerging as key players in multiple cancer types and thus represent an attractive target for pharmacological intervention (Kimes et al., 2014; Oltean and Bates, 2014; Wang and Lee, 2018; Zhang and Manley, 2013). To date however, the specific role of PGC-1αalternative splice variants in cancer has remained elusive. We demonstrate that the CRTC1-MAML2 fusion selectively induces CREB-dependent expression of the PGC-1α4 splice variant, which we and others have shown is associated with IGF-1 expression in anabolic skeletal muscle (Bruno et al., 2014; Ruas et al., 2012; Tasoulas et al., 2019). We confirm here that CRTC1-MAML2 directs similar anabolic pro-growth and pro-survival signaling by coordinating this autocrine PGC-1α4 – IGF-1 signal circuit in fusion positive salivary MEC.

A model that may explain the transcriptional and pre-mRNA splicing effects of CRTC1-MAML2 on PGC-1α4 is that this fusion acquires a strong transcriptional activation domain from MAML2 and retains the CREB binding domain of CRTC1, however the splicing domain normally present in full-length CRTC1 is deleted (Amelio et al., 2009; Tasoulas et al., 2019). Importantly, the PGC-1α4 variant retains the activation domain common to other PGC-1α transcript isoforms, which mediates binding to nuclear hormone receptor transcription factors such as PPARγ (Li et al., 2008; Martínez-Redondo et al., 2015; Sonoda et al., 2007). Similar to PGC-1α, dichotomous roles for PPARγ have been described with evidence pointing to oncogenic activity in some cancers but tumor suppressor actions in other cancers, and this is supported by data using PPARγ agonists or antagonists, respectively (Goldstein et al., 2017; Kardos et al., 2016; Khandekar et al., 2018; Zou et al., 2019). We demonstrate here that PGC-1α4 upregulates IGF-1 in a PPARγ-dependent manner in CRTC1-MAML2 positive salivary MEC, which sensitizes these tumor cells to treatment with PPARγ inverse agonists. Given the limitations of anti-IGF1R monotherapy, the identification of PPARγ inverse agonists as potential therapeutics for the treatment of MEC is particularly exciting. A limitation of this study, however, is that therapeutic effects achieved with the PPARγ inverse agonists SR10221 and SR2595 were in the low micromolar range, despite the fact that functional ligand binding and induced PPARγ conformational changes are possible in the nanomolar ranges (Marciano et al., 2015; Shang et al., 2020). These differences in the inverse agonist concentrations required to achieve corepressor-dependent PPARγ repression may be due to higher binding affinity of the PGC-1α4 variant with PPARγ and/or the increased levels of competing endogenous ligands (e.g. esterified lipids) in salivary MEC cells. Interestingly, our bioinformatic analyses revealed that several metabolic pathways with the potential to directly influence potential endogenous ligand levels, including lipid biosynthesis and modification pathways, are significantly elevated in CRTC1-MAML2 positive salivary MEC (Table S2).

In summary, our study identifies the CRTC1-MAML2 fusion oncogene as a master regulator of transcriptional networks that promote tumorigenic processes in part via a novel PPARγ-dependent PGC-1α4 – IGF-1 signaling circuit. Therefore, elucidation of the CRTC1-MAML2-regulated PGC-1α4-dependent transcriptional program may open new avenues for the identification of other pathways that can be exploited in order to stratify patients suitable for precision therapies. Our findings establish that targeting this ‘synthetic’ signaling circuit via IGF-1R inhibition or PPARγ inverse agonism, either individually or in combination, is a potential therapeutic option for patients with CRTC1-MAML2 positive salivary MEC.

## ACKNOWLEDGMENTS

The authors thank Gary Johnson and Brian Golitz for technical support in performing the small molecule drug screen. We would also like to thank Al Baldwin, Gaorav Gupta, Wendell Yarbrough, and members of the Amelio Lab for helpful discussions, suggestions, and/ or scientific review of this manuscript. The UNC Lineberger Animal Studies, Animal Histopathology, Small Animal Imaging, Flow Cytometry, Office of Genomics Research, Translational Genomics Lab, Translational Pathology Lab, High Throughput Sequencing Facility, and Bioinformatics Cores are all supported in part by an NCI Center Core Support Grant (P30-CA016086) to the UNC Lineberger Comprehensive Cancer Center. The UNC Flow Cytometry Core Facility is also supported by the North Carolina Biotech (NCBT) Center Institutional Support Grant 2012-IDG-1006. The Translational Pathology Lab is also supported by NIH U54-CA156733, NIEHS P30-EOS010126, and NCBT 2015-IDG-1007. This work was supported in part by a NIH/NIGMS T32-GM007092 and NIH/NIDCR F31-DE027282 training grant (to A.M. Musicant), NIH/NIGMS R01-GM130866 and the American Heart Association 19CDA34660248 (to J. G.), Head and Neck Cancer Fund (to T.G. Hackman and D.N. Hayes), UNC Translational Team Science Award (TTSA; to T.G. Hackman and A. L. Amelio), University Cancer Research Fund (UCRF; to A.L. Amelio), and by a NIH/NCI Howard Temin Pathway to Independence Award in Cancer Research R00-CA157954 (to A.L. Amelio).

## AUTHOR CONTRIBUTIONS

Conception and design, A.M.M., K. P-S., and A.L.A.; Development of methodology, A.M.M., K. P-S., W. G., A. C., C. A. F., D.N.H., and A.L.A.; Acquisition of data (provided animals, acquired and managed patient data, provided facilities, etc.), A.M.M., K. P-S., W. G., M. S., A. C., E. C. H., Y-H. T., M. C. H., S. S., R. B., T. G. H., R. P., C. A. F., and A.L.A.; Analysis and interpretation of data (e.g., statistical analysis, biostatistics, computational analysis), A.M.M., K. P-S., Y-H. T., J. S. P., J. G., C. A. F., D.N.H., and A.L.A..; Writing of the manuscript, A.M.M. and A.L.A.; Review and/or revision of the manuscript: A.M.M., K. P-S., W. G., Y-H. T., T. G. H., J. S. P., J. G., C. A. F., D.N.H., and A.L.A.; Administrative, technical, or material support (i.e., reporting or organizing data, constructing databases), A. C., E.C.H., M. C. H., R. B., and J.S.P.; Study supervision: J.S.P., C.A.F., D.N.H., and A.L.A.

---

**Supplemental Figure S1. Bioinformatics analyses reveal that IGF-PI3K signaling is deregulated in C1/M2 fusion positive salivary MEC tumors, Related to Figure 1.**

A. Unsupervised heat map of the top 1500 most variable genes between MEC and normal salivary gland samples. Normal (e.g. N #1, N #2) and tumor (e.g. Case #1, Case #7) samples are indicated at the top of the heat map in black and green, respectively. C1/M2 fusion status is indicated in blue and grey.
B. Gene set enrichment analysis (GSEA) of IGF-1 pathway genes. Circle sizes in the bubble plot indicate the proportion of genes within each pathway that are statistically significant (adjusted p-value <0.05). Grey bubbles indicate q-value ≥0.05 and red bubbles indicate q-value <0.05.
C. *left*, Gene set enrichment plot for genes in the GSEA Molecular Signatures Database (MSigDB) GNF2_IGF1 dataset in fusion-positive MEC and normal salivary gland samples. *right*, Violin plot showing mean VST counts of genes in the GNF2_IGF1 gene list in fusion-positive MEC, fusion-negative MEC, and normal salivary gland samples.
D. Gene set enrichment plot for IGF1-related genes in our curated Musicant_MEC_CRTC1-MAML2_IGF1 gene set in fusion-positive MEC and normal salivary gland samples.

**Supplemental Figure S2. Validation of IGF-1 expression in C1/M2 fusion positive salivary MEC cell lines compared to fusion negative cell lines, Related to Figure 2.**

A. Comparison of *C1/M2* and *IGF-1* CT values in qPCR data from cell lines. C1/M2-negative cell lines are indicated in grey and C1/M2-positive cell lines are indicated in red. The fitted regression (solid) line demonstrates that the expression levels of the two genes are correlated (r^2^ = 0.8664, p = 0.0008). 95% confidence intervals are indicated by curved dashed lines. Gene expression data is the average of at least 3 independent experiments.
B. Histologic and immunohistochemical analysis of C1/M2-positive and C1/M2-negative xenografts. Representative xenograft sections were formalin fixed, embedded, and H&E stained (top; magnification: 200x; scale bar: 100 µm), revealing epidermoid cells in C1/M2-negative xenografts and epidermoid, mucus, and intermediate cells in the C1/M2-positive xenografts. Mucicarmine staining (middle; magnification: 400x; scale bar: 100 µm) revealed mucus cells (indicated by yellow arrows) in C1/M2-positive xenografts. IHC staining for IGF-1 (bottom; magnification: 200x; scale bar: 100 µm) demonstrated that IGF-1 expression is increased in C1/M2-positive xenografts relative to C1/M2-negative xenografts.

**Supplemental Figure S3. Small molecule drug screen overview, Related to Figure 2.**

A. Analysis of drug screen robustness based on a Z’-factor analysis. Data points in red indicate viability of cells treated with positive control (1 µM Bortezomib-100% cell death) and data points in blue indicate viability of cells treated with negative control (DMSO-0% cell death). Cell lines are listed along the x-axis, with fusion status (C1/M2-positive or - negative) indicated below in grey (negative) or red (positive). Z’-factor >0.75 confirms a robust dynamic range of drug effects indicating good drug screen reliability.
B. Waterfall plots depicting the effect of individual candidate drugs in C1/M2-positive cell lines (colored points) relative to the effect of the same drug in the C1/M2-negative control cell line (A253). Points above the dashed line indicate that the tested drug is more effective in the C1/M2-positive line, while points below the dashed line indicate that the tested drug is more effective in A253 cells. Several IGF-1R inhibitors (BMS-754807, NVP-AEW541, and GSK1838705A) are more effective in C1/M2-positive cells lines (>50%); the relative effects of these inhibitors in each cell line are denoted with the indicated symbols.
C. Relative viability scores for each individual drug per cell line are shown as heatmaps. Scores were normalized to the respective DMSO conditions for each cell line, and drugs were ordered based on their efficacy against A253 fusion negative salivary epidermoid carcinoma cells. 0= no effect relative to DMSO, 1.0= 100% cell death, and a negative score= increased growth.

**Supplemental Figure S4. Drug screen validation of IGF-1R inhibition by genetic and pharmacologic approaches, Related to Figure 3.**

A. Quantitative real-time PCR analysis of *IGF-1R* mRNA levels in cells transduced with lentiviruses expressing either a non-specific shRNA (shNS) or two different shRNAs targeting IGF-1R (shIGF-1R_#1 and shIGF-1R_#2). Relative fold expression is shown normalized to *RPL23* mRNA levels (n = 2, mean ± SEM). \*\**p* ≤ 0.01.
B. Western blot analysis demonstrating relative knockdown of IGF-1Rβ protein levels with shIGF1R_#1 and shIGF1R_#2 relative to control shRNA (shNS).
C. Number of colonies formed by HMC1 (left), HMC3A (center), and HMC3B (right) cells treated with DMSO or BMS-754807 at ½ IC50 concentration or IC50 concentration (*see Table 1 for IC50 values*). A colony is defined as a cluster of ≥50 cells (n = 3 biological replicates, mean ± SEM). \**p* ≤ 0.05, \*\**p* ≤ 0.01.

**Supplemental Figure S5. C1/M2 Regulates IGF-1 Transcription Through PGC1α4-PPARγ, Related to Figure 4**

A. Quantitative real-time PCR analysis of *C1/M2* mRNA levels in C1/M2-inducible cells (doxycycline-regulated HEK293-*CMV^TetR^TetO^C1/M2^* cells) treated with and without 1 µg/mL doxycycline. Relative fold expression is shown normalized to *RPL23* mRNA levels. (n = 3 biological replicates, mean ± SEM). ****p < 0.0001.
B. Western blotting analysis of IGF-1 and tubulin protein expression levels in HEK293-*CMV^TetR^TetO^C1/M2^* cells treated with and without 1 µg/mL doxycycline.
C. *left*, Schematic depicting the AlphaLISA assay. Donor and acceptor beads each carry an antibody that detects the analyte of interest. Donor beads can be excited by light at 680 nm and can then convert molecular oxygen to its excited state. Acceptor beads can utilize the excited oxygen molecule to emit light at 615 nm. The transfer of this excited molecular oxygen can only occur when donor and acceptor beads are brought into close proximity through mutual binding of the analyte. Thus, quantification of light at 615 nm allows for the indirect quantification of analyte levels. *right*, Standard curve showing the relationship between AlphaLISA signal and IGF-1 protein concentration.
D. Quantitative real-time PCR analysis of *C1/M2* (*left*) and *NR4A2* (*right*) mRNA levels in HMC3A cells transduced with a non-specific shRNA (shNS) or two different shRNAs targeting C1/M2 (shMAML2_#1 and shMAML2_#2). Relative fold expression is shown normalized to *RPL23* mRNA levels. (n = 3 biological replicates, mean ± SEM). ns = not significant, \**p* ≤ 0.05, \*\**p* ≤ 0.01.
E. Quantitative real-time PCR analysis of *PGC-1α4* mRNA levels in HMC3A-*PGK^TetOn3G^TRE3GS^dnCREB^* cells treated with and without 1 µg/mL doxycycline. Relative fold expression is shown normalized to *RPL23* mRNA levels (n = 3 biological replicates, mean ± SEM). ***p < 0.001.
F. *left*, Schematic depicting stable HEK293-*PGK^TetOn3G^TRE3GS^PGC-1α4^* cells engineered to inducibly express *Flag-PGC-1α4* upon treatment of cells with tetracycline/doxycycline. *right*, Quantitative real-time PCR validation of *PGC-1α4* following 24 hr treatment with increasing concentrations of doxycycline in HEK293-*PGK^TetOn3G^TRE3GS^PGC-1α4^* cells. (n = 3 biological replicates, mean ± SEM). ****p < 0.0001. HEK^WT^ = non-transduced HEK293A cells.
G. Western blot analysis of Flag-PGC-1α4 and tubulin protein expression levels in HEK293-*PGK^TetOn3G^TRE3GS^PGC-1α4^* cells treated with increasing concentrations of doxycycline for 24 hr. Blot is representative of two biological replicates.
H. Cell proliferation assay showing relative confluency of HMC3A cells with and without shRNA-mediated knockdown of *PGC-1α* (*left*), *IGF-1R* (*center*), or *C1/M2* (*right*). Time to 50% confluence is shown by dashed lines. Data shown is representative of three independent experiments (n = 3 biological replicates). shNS = non-specific shRNA; shPGC-1α_#1 and _#2 = shRNAs targeting *PGC-1α*; shIGF-1R_#1 and _#2 = shRNAs targeting *IGF-1R*; shMAML2_#3 = shRNA targeting *C1/M2* and *MAML2*.

**Supplemental Figure S6. Validation of PPARγ-dependent IGF-1 transcriptional activation by a C1/M2-mediated PGC1α4 circuit, Related to Figure 5.**

A. Schematic of the human IGF-1 gene locus, highlighting the P2 promoter (located upstream of the second exon) as well as several PPARγ binding motifs (PPARG motifs) located within this promoter region.
B. Light output generated by an IGF-1 P2 promoter-driven luciferase reporter in C1/M2-inducible cells (doxycycline-regulated HEK293-*CMV^TetR^TetO^C1/M2^* cells) treated with and without 1 µg/mL doxycycline treatment (n = 3 biological replicates, mean ± SEM). \**p* ≤ 0.05, \*\**p* ≤ 0.01, ***p < 0.001.
C. Light output generated by a PPARγ-response element (PPRE)-driven luciferase reporter in HEK293-*CMV^TetR^TetO^C1/M2^* cells with and without 1 µg/mL doxycycline treatment (n = 3 biological replicates, mean ± SEM). \**p* ≤ 0.05.
D. PPARγ-response element (PPRE)-driven (*left*) and IGF-1 promoter-driven (*right*) luciferase reporter assay showing activation of basal activity of endogenously expressed PPARγ in HEK293-*CMV^TetR^TetO^C1/M2^* cells. Cells were treated with increasing concentrations of the PPARγ agonist GW1929 for 24 hr prior to measuring luciferase activity (n = 2 biological replicates, mean ± SEM).
E. Number of colonies formed by HMC3A and HMC3B cells treated with DMSO or SR10221 at ½ IC50 concentration or IC50 concentration (*see Table 2 for IC50 values*). A colony is defined as a cluster of ≥50 cells (n = 3 biological replicates, mean ± SEM). ND = not detected, \**p* ≤ 0.05, \*\*\**p* ≤ 0.001.

## STAR METHODS

### LEAD CONTACT AND MATERIALS AVAILABILITY

Further information and requests for resources and reagents should be directed to and will be fulfilled by the Lead Contact, Antonio L. Amelio. The plasmids have been deposited with Addgene and cell lines generated in this study are available upon request via a material transfer agreement (MTA).

### EXPERIMENTAL MODEL AND SUBJECT DETAILS

#### Clinical samples

All research involving human tumor tissues was reviewed and approved by The University of North Carolina at Chapel Hill Institutional Review Board under IRB protocols 15-1604 and 17-2947. Tissues from these de-identified clinical subjects were identified from chart review and archived FFPE salivary MEC (*n* = 20) or normal salivary gland (*n* = 6) tissue samples stored at room temperature less than ten years before blocks were sectioned and RNA isolation performed. For all cases, multiple H&E slides were reviewed from each case by a pathologist and sections with tumor were selected for inclusion in the study. Adjacent serial unstained sections were then macrodissected and tumor material submitted for RNA extraction.

#### Xenograft models

All animal studies were reviewed and approved by The University of North Carolina at Chapel Hill Institutional Animal Care and Use Committee under IACUC protocol 17-202. Male 6-8 week old athymic nude mice (Nu/Nu) were obtained from the Animal Studies Core at the University of North Carolina at Chapel Hill and housed in facilities run by the Division of Comparative Medicine at the University of North Carolina (Chapel Hill, NC, USA). For all xenograft studies, mice were subcutaneously injected with 1×10^6^ UM-HMC-3A cells resuspended in 50% HBSS and 50% Matrigel. When the average tumor size reached a palpable size (∼50 mm^3^), animals were randomized into two groups (*Vehicle* and *SR10221* or *SR2595*) so that the average tumor size in each group was approximately equal. The vehicle group was intraperitoneally (i.p.) injected daily with 250 uL of a 10% DMSO, 10% Tween-80, 80% PBS solution. Drug-treated animals were i.p. injected daily with a 10% DMSO, 10% Tween-80, 80% PBS solution containing either 20 mg/kg SR10221 or 60 mg/kg SR2595 (Sigma-Aldrich #SML2037) based on an average mouse weight of 25g. Caliper measurements were collected every 2 days and bioluminescent imaging (BLI) was performed every 5 days throughout the course of the study.

#### Cell lines

UM-HMC-1 (HMC1), UM-HMC-3A (HMC3A), and UM-HMC-3B (HMC3B) cells (Warner et al., 2013) were kindly provided by Dr. Jacques Nör (University of Michigan, Ann Arbor, MI, USA). NCI-H292 (H292) and NCI-H3118 (H3118) cells (Tonon et al., 2003) were generously provided by Dr. Frederic Kaye (University of Florida, Gainesville, FL, USA). A253, A388, and A431 cells were kindly provided by Dr. Bernard Weissman (University of North Carolina, Chapel Hill, NC, USA). HMC1 (source: male, salivary gland mucoepidermoid carcinoma (MEC)), HMC3A (source: female, hard palate MEC), and HMC3B (source: female, lymph node metastasis of hard palate MEC) parental and stably transduced lines were cultured in DMEM medium (Gibco #11965-118) supplemented with 10% FBS (Atlanta Biologicals #S11550), 20 ng/mL EGF (Sigma-Aldrich #E9644), 400 ng/mL hydrocortisone (Sigma-Aldrich #H0888), 5 μg/mL insulin (Sigma-Aldrich #I6634), and 1X pen/strep/glutamine (PSG; Life Tech #10378016). H292 (source: female, lung MEC) cells were cultured in RPMI-1640 medium (Life Tech #11875119) supplemented with 10% FBS, 1X GlutaMAX (Life Tech #35050061), 1X NEAA (Gibco #11140050), 1X NaPyr (Life Tech #11360070), and 1X PSG. HEK293A, HEK293-*CMV^TetR^TetO^C1/M2^* (Amelio et al., 2014), HEK293-*PGK^TetOn3G^TRE3GS^PGC-1α4^*, Lenti X-293T (viral packaging cell line; Takara #632180), H3118 (source: female, parotid gland MEC), A388 (source: male, squamous cell carcinoma (SCC)), and A431 (source: female, vulvar SCC) cells were cultured in DMEM supplemented with 10% FBS, 1X GlutaMAX, and 1X PSG. A253 (source: male, salivary gland SCC) cells were cultured in McCoy’s 5A medium (Life Tech #16600108) supplemented with 10% FBS, 1X GlutaMAX, and 1X PSG. Cells were passaged using TrypLE (Gibco #12604013) every 2-3 days or when they reached 90% confluence. All cells were maintained in a 37°C, 5% CO2 atmosphere. All cell lines were confirmed mycoplasma-free by PCR as previously described (Young et al., 2010) and using mycoplasma detection primers (see Table S6).

### METHOD DETAILS

#### Data and Code Availability

All gene expression data for this study are available at the Gene Expression Omnibus under GSEXXXX. Sequence data from which the gene expression analyses were derived are available at the Short Read Archive under SRPXXXX with authorization controlled by dbGaP.

#### Clinical RNA isolation

Formalin-fixed paraffin-embedded (FFPE) tissue samples were sent to the UNC Lineberger Comprehensive Cancer Center Translational Genomics Lab (TGL) for RNA isolation using the Maxwell 16 MDx Instrument (Promega #AS3000) and the Maxwell 16 LEV RNA FFPE Kit (Promega #AS1260) according to the manufacturer’s protocol (Promega #9FB167). Pathology review of a hematoxylin and eosin (H&E) stained slide was used to guide macro-dissection of unstained slides to enrich for tumor RNA. Total RNA quality was measured using a NanoDrop spectrophotometer (Thermo Scientific ND-2000C) and a TapeStation 4200 (Agilent G2991AA). Total RNA concentration was quantified using a Qubit 3.0 fluorometer (Life Technologies Q33216).

#### RNA-seq

Total RNA sequencing libraries were prepared at TGL using a Bravo Automated Liquid-Handling Platform (Agilent G5562A) and the TruSeq Stranded Total RNA Library Prep Gold Kit (Illumina 20020599) according to the manufacturer’s protocol (Illumina 1000000040499). RNAseq library quality and quantity were measured using a TapeStation 4200 (Agilent G2991AA), pooled at equal molar ratios, and denatured according to the manufacturer’s protocol (Illumina 15050107). Sequencing was performed at the High Throughput Sequencing Facility (HTSF) at UNC Chapel Hill. Two RNA-seq libraries were sequenced per lane on a HiSeq2500 (Illumina SY–401–2501) with 2×50 bp paired-end configuration according to the manufacturer’s protocol (Illumina 15035786).

#### Bioinformatics

RNAseq data analyses were performed with FASTQ files aligned to the GRCh38 human genome (GRCh38.d1.vd1.fa) using STAR v2.4.2 (Dobin et al., 2013) with the following parameters: -- outSAMtype BAM Unsorted, --quantMode TranscriptomeSAM. Transcript abundance for each sample was estimated using Salmon version 0.1.19 (Patro et al., 2017) to quantify the transcriptome defined by Gencode v22. Gene level counts were summed across isoforms and genes with low expression (i.e. samples with fewer than 10 reads) were removed prior to downstream analyses. The R package DESeq2 (version 1.24.0) was used to test for differentially expressed genes between CRTC1-MAML2 positive MEC tumors and normal samples or between all MEC tumors and normal samples. The C1/M2-regulated IGF-PI3K gene signature (Musicant_MEC_CRTC1-MAML2_IGF1 curated gene set) includes genes that are differentially expressed between CRTC1-MAML2 positive MEC samples and normal samples with Benjamini– Hochberg FDR (q-value) < 0.05 and a fold change > 2. The resulting list of genes was refined to include only genes shared with IGF1-related Molecular Signatures Database (MSigDB) curated gene sets. Gene set enrichment analysis was performed on a customized list of gene sets with GSEA software from the Broad Institute (Subramanian et al., 2005). The customized gene sets include a list of pathways from MSigDB curated gene sets that are related to IGF1. Hierarchical clustering and heat-map plotting were performed using the ComplexHeatmap R package version 2.0.0 (Gu et al., 2016). An average linkage algorithm with a Euclidean distance function was applied to variance stabilizing transformed (VST) gene expression data for the gene signature.

#### RTKi drug screen

Compound screening was performed as previously described (Bevill et al., 2019; Lipner et al., 2020). Briefly, drug screening was performed in a 384 well plate format, in duplicate. Cells were seeded in 45 µL of full culture media; A253 (1000 cells/well), H292 (600 cells/well) H3118 (7000 cells/well), HMC1 (500 cells/well), HMC3A (500 cells/well) and HMC3B (700 cells/well). Cells were seeded in 384-well plates using a BioTek microplate dispenser. Cells were allowed to adhere overnight, following which they were treated with 176 individual drugs at 6 doses (10 nM, 100 nM, 300 nM, 1 µM, 3 µM and 10 µM) using a Beckman Coulter Biomek FX Automated Liquid Handling instrument. Each plate also included 1 µM Bortezomib and 1% DMSO as a positive and negative controls for growth inhibition, respectively. Furthermore, a full 384 well plate with 1% DMSO treatment was also seeded per cell line and was used to confirm minimal well-location associated intra-plate variability. 72 hr post drug addition, cells were lyzed and cell viability was measured using Cell Titer Glo 2.0 (Promega, #G9243), according to the manufacturer’s instructions. Luminescence was measured using a PHERAstar FS instrument and growth inhibition was calculated relative to DMSO-treated wells. Z’ scores were calculated as previously described (Zhang et al., 1999). The 1 µM Bortezomib (positive; 0% viability) and 1% DMSO (negative; 100% viability) were used to calculate the dynamic range and screen variability/reliability (Z’ score). A Z’ score of >0.75 was obtained for all cell lines tested. The 3SD/5SD cutoffs were calculated based on the variability observed in the 1% DMSO-negative control plates/wells.

#### Virus production and transduction

Lentiviral or retroviral expression plasmids were co-transfected with VSV-G envelope plasmid and either δ8.2 gag/pol (lentivirus) or pMD (retrovirus) helper plasmids into Lenti X-293T cells seeded in 10 cm tissue culture dishes. Transfections were performed using 1 mg/mL Polyethyleneimine (PEI) Transfection Reagent (VWR #BT129700). Briefly, 1.5 μg VSV-G, 5 μg δ8.2 or pMD, and 6 μg lentiviral or retroviral plasmid were brought to 500 μL with OptiMEM (LifeTech #1158021) and vortexed briefly. In a separate tube, 25 μL PEI (2 µL PEI/µg DNA) was added to 475 μL OptiMEM and vortexed briefly. Both solutions were incubated at room temperature for 5 min, then combined and incubated for an additional 20 min at room temperature. This mixture was then added dropwise to the seeded Lenti X-293T cells. The next day, cell culture media was replaced with DMEM supplemented with 1x NaPyr, 10 mM HEPES, 1X GlutaMAX, and 1X PSG (no FBS). Two days later, media was collected and filtered through a 0.45 μm PVDF membrane and then viral particles were concentrated via ultracentrifugation (100,000g for 2 hr) into a sucrose cushion. Concentrated virus was resuspended in cold PBS and either stored at −80°C or used immediately for transduction.

Stable cells were generated by spinfection of various cell lines seeded at 50% confluency in 12-well plates using concentrated virus and 4 µg/mL polybrene added directly to cells prior to centrifugation at 1200g for 90 min at 30°C. To generate stable LumiFluor-overexpressing MEC cells, HMC3A cells were transduced with concentrated lentivirus expressing our GpNLuc reporter (Schaub et al., 2015) and selected with puromycin for at least 10 days prior to use in experiments. To generate shRNA-expressing cells, parental HMC3A or HMC3A-*PGK^GpNLuc^* cells were transduced with concentrated shRNA-expressing retro- or lentivirus expressing the shRNA of interest and selected with geneticin or puromycin, respectively, for at least 10 days prior to use in experiments. Proliferation assays, quantitative real-time PCR, and Western blotting were performed on polyclonal cell populations within 14 days of transduction for all shRNA knockdown experiments. To generate cells that express the PGC-1α4 splice variant, HEK293A cells were first transduced with a concentrated lentivirus that expresses TetOn3G and stable cells selected with puromycin. Subsequently, these HEK293-*PGK^TetOn3G^* stables were transduced with a concentrated lentivirus that expresses PGC-1α4 upon tetracycline/doxycycline administration and the resulting HEK293-*PGKTetOn3G-TRE3GS^PGC-1α^*^4^ stable cells selected with hygromycin for at least 10 days prior to use in experiments. To generate MEC cells with inducible dominant-negative CREB (dnCREB; A-CREB), HMC3A cells were first transduced with a concentrated lentivirus that expresses TetOn3G and stable cells selected with puromycin. These HMC3A-*PGK^TetOn3G^* stables were transduced with a concentrated retrovirus that expresses A-CREB (Ahn et al., 1998) upon tetracycline/doxycycline administration, and the resulting HMC3A-*PGK^TetOn3G^-TetO^dnCREB^* stable cells selected by fluorescence activated cell sorting (FACS) by gating for tdTomato-positive cells.

#### Cell viability

HMC1, UHMC3A, or HMC3B cells were seeded in 96-well clear-bottom, white-walled plates (Corning #3917) at 15,000 cells/well. The next day, media was replaced with 200 µL fresh media containing either vehicle (DMSO) or increasing concentrations of drug (BMS-754807 (Sigma-Aldrich #BM0003), PPP (Santa Cruz #SC-204008), SR10221, SR2595 (Sigma-Aldrich #SML2037), or T0070907 (Fisher #NC1015539); final concentration 1% DMSO). 72 hr later, the amount of ATP (a proxy for cell viability) in each well was measured using the ATPlite Luminescence Assay System (PerkinElmer #6016949) according to the manufacturer’s instructions. Briefly, plates were removed from the cell culture incubator and equilibrated for 30 min at room temperature in the dark. Then, media was aspirated from each well and 100 µL reconstituted ATPlite 1-step reagent was added to each well. Plates were shaken for 2 min (425 cpm, 3 mm orbit) in a Cytation 5 plate reader (BioTek Instruments, Inc.) and then ATP levels were quantified by measuring total luminescence. IC50 values were calculated in GraphPad Prism (GraphPad, San Diego, CA, USA) using a non-linear curve fit (log[agonist] vs response, four parameter-variable slope). All assays were performed in biologic triplicate with three technical replicates for each condition.

#### Cell proliferation assays (Cytation 5)

HMC3A-*PGK^GpNLuc^* cells (either non-transduced or transduced with shRNAs targeting PGC-1α, IGF-1R, or MAML2) were seeded in 12-well plates at 200,000 cells/well. The next day, fresh media was applied to each well and then individual wells were imaged at 0, 4, 8, 24, 28, 32, and 48 hr following the media change. Five separate fields of view were imaged for each well at 4X magnification using a fluorescent plate reader (BioTek Cytation 5, BioTek Instruments, Inc.). Between each imaging time point, plates were returned to the cell culture incubator. To determine cell confluence, a primary mask was applied to each fluorescent image using the recommended parameters set out in the BioTek Technical Note “Measuring Confluence Using High Contrast Brightfield” (https://www.biotek.com/resources/technical-notes/measuring-confluence-using-high-contrast-brightfield/). Confluence measurements from multiple fields of view in a single well were used as technical replicates. Each assay was performed in biologic triplicate.

#### Cell proliferation assays (IncuCyte Zoom)

HMC3A cells (either non-transduced or transduced with an shRNA targeting IGF-1R) were seeded in 48-well plates at 15,000 cells/well. The next day, media was replaced with fresh media containing either vehicle (1% DMSO) or the indicated concentration of PPP (Santa Cruz #SC-204008) or SR10221. Then, plates were imaged at 10X magnification every 2-4 hours on an IncuCyte Zoom (Essen BioScience), which maintained plates at a constant 37°C and 5% CO2 for the duration of the assay. Four separate fields of view were imaged for each well and an image mask was applied to each image to generate confluency measurements. Each assay was performed in biologic triplicate with at least three technical replicates per experiment.

#### Western blotting

Whole cell lysates were prepared in buffer containing 5% glycerol (Sigma-Aldrich #G5516), 25 mM Tris (pH 7.4; Fisher #BMA51237), 150 mM NaCl (Fisher #BMA51202), 1 mM EDTA (Sigma-Aldrich #E7889), and 1% NP-40 (Fisher #50-147-289) supplemented with protease inhibitors (cOmplete, EDTA-free Protease Inhibitor Cocktail, Roche #04693132001) and phosphatase inhibitors (PhosSTOP, Roche #10917400). Lysates (30-50 mg) were loaded onto mini-10% tris-glycine polyacrylamide gels and proteins were separated using sodium dodecyl sulfate-polyacrylamide gel electrophoresis (SDS-PAGE). Proteins were transferred to a 0.45 µm nitrocellulose membrane (Fisher #0088018) using a Bio-Rad Trans-Blot Turbo system set at 1.3 Amps (constant) and 25 V for 30 min. Membranes were blocked for 1 hr at room temperature in TBS-T + 5% milk and then incubated over night at 4°C with primary antibodies: IGF-1 (0.2 µg/mL; Abcam #ab9572), IGF-1Rβ (1:2500; Cell Signaling #3018S), actin (1:5000; Sigma #A3854), tubulin (1:5000; Sigma #T6074), or Flag (1:5000; Sigma #A8592) diluted in TBS-T + 5% milk. Following primary antibody incubation, membranes were washed and probed with Horseradish peroxidase (HRP)-conjugated goat anti-mouse (1:5000; Thermo Fisher #31432) or donkey anti-rabbit (1:5000; Thermo Fisher #31458) secondary antibodies diluted in TBS-T supplemented with 5% milk for 1-2 hr at room temperature. Blots were imaged using Clarity ECL (Bio-Rad #170-5060) and ImageQuant LHS4000 (GE).

#### 2D colony formation

HMC1, HMC3A, and HMC3B cells were seeded in 24-well plates at 800, 600, and 1400 cells/well, respectively. The next day, media was replaced with 500 µL fresh media containing either vehicle (DMSO) or the indicated concentration of drug (BMS-754807 (Sigma-Aldrich #BM0003), PPP (Santa Cruz #SC-204008), SR10221, SR2595 (Sigma-Aldrich #SML2037), or T0070907 (Fisher #NC1015539); final concentration 1% DMSO). Seven days later, media was removed from all wells and cells were fixed in 10% buffered formalin (Fisher #SF994) for 5 min at room temperature. Wells were washed once in ddH2O and then stained in 0.05% crystal violet (Sigma-Aldrich #C6158) for 30 min at room temperature. Wells were then washed an additional 3 times in ddH2O to remove any unbound stain and allowed to dry at room temperature overnight. Once dry, individual wells were imaged at 4X magnification on a BioTek Cytation 5 plate reader (BioTek Instruments, Inc.) and images were stitched using the Gen5 software (v2.09; BioTek Instruments, Inc.) using the default parameters. Colony numbers were quantified manually in Image J, where a colony is defined as a cluster of at least 50 individual cells.

#### 3D tumor spheroid formation

Individual wells of a 48-well plate were coated with 100 µL of a 25% Matrigel (Fisher #CB4023A) solution diluted in cold 1X DPBS. The plate was spun in a Thermo Scientific Sorvall LYNX 4000 centrifuge at 4°C for 10 min at 2000g to ensure even distribution across the well surface. The plate was then placed in a 37°C incubator for 1 hr to allow the Matrigel ‘bed’ to completely polymerize. Next, excess DPBS was gently aspirated from the wells and 2000 HMC3A cells resuspended in 100 µL media were added to each well. The plate was centrifuged at 4°C for 5 min at 2000g to embed the cells into the Matrigel. The plate was returned to the incubator for an additional hour to allow for cell adhesion to the Matrigel. After 1 hr, excess media was carefully aspirated and 50 µL undiluted Matrigel was gently added to each well on top of the cells. The plate was again returned to the incubator for 1 hr to allow for Matrigel polymerization. Finally, 500 µL media (containing either drug or vehicle at a final concentration of 1% DMSO) was added to each well. Every 24 hr, wells were imaged at 4X magnification on a BioTek Cytation 5 plate reader (BioTek Instruments, Inc.) and images were stitched using the Gen5 software (v2.09; BioTek Instruments, Inc.) using the default parameters. Tumorsphere area was manually measured in ImageJ using the ROI area measurement tool. A minimum of 50 individual tumorspheres were quantified for each condition in each experiment.

#### Caspase 3/7 apoptosis assay

HMC3A cells were seeded in 6-well plates at 350,000 cells/well. The next day, media was replaced with 2 mL fresh media containing either vehicle (DMSO) or the indicated concentration of drug PPP (Santa Cruz #SC-204008) or SR10221; final concentration 1% DMSO). At the indicated time point, cells were trypsinized, spun down, and resuspended in 1 mL 1X DPBS supplemented with 1X GlutaMAX (Life Tech #35050061) and 10% FBS (Atlanta Biologicals #S11550). Apoptotic cells were visualized using the CellEvent Caspase 3/7 Green Flow Cytometry Assay Kit (ThermoFisher #C10427) according to the manufacturer’s instructions. Briefly, 1 µL CellEvent Caspase 3/7 Green Detection Reagent was added per 1 mL of resuspended cells. This mixture was gently vortexed and incubated at 37°C for 25 min. Next, 1µL SYTOX AADvanced Dead Cell Stain was added per 1 mL of resuspended cells. This mixture was gently vortexed and incubated at 37°C for an additional 5 min. Samples were analyzed via flow cytometry on a BD Accuri C6 instrument. An aliquot of unstained cells from each condition was reserved as a control to generate gates.

#### Chromatin immunoprecipitation (ChIP) assay

HEK293-*CMV^TetR^TetO^C1/M2^* cells (Amelio et al., 2014) were seeded in 15 cm tissue culture dishes in either plain growth medium or growth medium containing 1 µg/mL doxycycline (Sigma-Aldrich #D9891). After 48 hr, formaldehyde (Fisher #BP531) was added dropwise to each plate to 1% final concentration. Plates were incubated at room temperature for 10 min and then crosslinking was stopped by adding glycine (Fisher #BP381-1) to a final concentration of 120 mM and incubating for an additional 5 min. Cells were then transferred to a tube, pelleted at 4°C, and washed twice with cold 1X DPBS. Finally, washed cells were resuspended in 1X DPBS supplemented with protease inhibitors (cOmplete, EDTA-free Protease Inhibitor Cocktail, Roche #04693132001) to a concentration of 5×10^6^ cells/mL and stored at −80°C in aliquots of 1 mL until lysis. For lysis, individual cell aliquots were pelleted, resuspended in 200 uL SDS lysis buffer (1% SDS, 10 mM EDTA, 50 mM Tris-HCl, pH 8.1) supplemented with protease inhibitors, and rocked at 4°C for at least 20 min. Next, 800 uL ChIP dilution buffer was added and aliquots were immersed in a 100% ethanol ice bath (−11 to −7°C) and sheared (2 Amp for 5 sec) using a sonicator (Qsonica #Q700A) equipped with a microtip sonic dismembrator (Model 505, Fisher #4418). A fraction of this sheared sample was reserved as a Pre-IP control. To the rest of the sample, 1 µg anti-Flag M2 antibody (Sigma #F1804) was added and samples were rocked at 4°C overnight. Antibody-chromatin complexes were isolated by rocking with Protein G SureBeads (BioRad #1614821) at 4°C for 4 hr. Beads were washed once each in low salt buffer (0.1% SDS, 1% TritonX-100, 2mM EDTA, 20 mM Tris-HCl, 15 mM NaCl), high salt buffer (0.1% SDS, 1% TritonX-100, 2 mM EDTA, 20 mM Tris-HCl, 500 mM NaCl), and LiCl buffer (250 mM LiCl, 1% NP-40, 1% Deoxycholate, 1mM EDTA, 10 mM Tris-HCl, pH 8.1) and then twice in TE buffer (1.2 mM EDTA, 10 mM Tris-HCl, pH 8.1). Samples were eluted twice with pre-heated (55°C) elution buffer for 15 min at room temperature with shaking. Cross-links were reversed by adding NaCl to 200 mM and incubating for at least 4 hr at 65°C. Samples were treated for 30 min at 37°C with RNase A at 8 µg/µL final concentration. Finally, samples were treated with Proteinase K solution (10 µL 0.5M EDTA, 10 µL 1M Tris, 1 µL 20 mg/mL Proteinase K (LifeTech #AM2546) for a 500 µL sample) for 1 hr at 50°C and then DNA was purified using the NucleoSpin Gel and PCR Cleanup Kit (Macherey-Nagel #740609.250). Occupancy of target proteins of interest within the promoter regions was assessed by qPCR (Table S6).

#### Luciferase reporter assay

HEK293-*CMV^TetR^TetO^C1/M2^* cells were seeded in 24-well plates at 100,000 cells/well. The next day, transfected each well with 500 ng of either PGC-1αproximal promoter or PGC-1α distal promoter driven luciferase reporter using Lipofectamine (Invitrogen #50470) according to the manufacturer’s protocol and using a DNA:lipid ratio of 1:2. The next day, media was replaced with 1 mL fresh media with or without 1 µg/mL doxycycline (Sigma-Aldrich #D9891). After 24 hr, media was removed from all wells and 100 μL 1% Triton X-100 was added to each well. Plates were rocked for 15 min at room temperature. 50 μL of the supernatant from each well was transferred to a white, opaque 96-well plate, and 50 μL Bright-Glo Luciferase Assay Buffer (Promega #E264B) was added to each well. Total luminescence was quantified immediately on a BioTek Cytation 5 plate reader (BioTek Instruments, Inc.). For the dose-response luciferase reporter assays, HEK293 cells were reverse transfected with 3xPPRE promoter or IGF1-P2 promoter driven luciferase reporters using Lipofectamine 2000 Transfection Reagent (Invitrogen #11668019) in 96-well plates at a density of 2X10^4^ cells/well. Forty-eight hours post-transfection cells were treated with either vehicle (DMSO) or the PPAR ligands GW1929, SR2595 or SR10221 at indicated concentrations. 24 hr after treatment, cells were lysed, and luciferase activity was quantified using the Dual-Glo Luciferase Assay System (Promega #E2920). Luminescence was measured using Synergy Neo microplate reader (BioTek). Values were normalized using Renilla expression.

#### Bioluminescence imaging

Bioluminescent-fluorescent BRET signal was measured non-invasively as previously described (Schaub et al., 2015) with minor modification. Briefly, animals were i.p. injected with 250 μM (1:20 dilution, ∼500 μg/kg) Nano-Glo Luciferase Assay Substrate (Promega, #N1120) in sterile PBS. Isoflurane-anesthetized animals were then imaged using an AMI Optical Imaging System (Spectral Instruments Imaging, Inc.) 5 min after injection. Images were captured with open filter and acquisition times of 5 min or less at the indicated settings. Data were analyzed using Aura imaging software (v2.2.0.0).

#### Histology and immunohistochemistry

All animals showing obvious tumors or other signs of distress were euthanized and subjected to full necropsy. For histological analysis, all tissues (including submandibular glands, sublingual glands, parotid glands, pancreas, and lungs) were fixed in 10% neutral buffered formalin for approximately 1 week at room temperature. Following fixation, tissues were processed on an ASP6025 automated tissue processor (Leica Biosystems) and embedded in paraffin wax. Blocks were sectioned at 4-6 μm, mounted on glass slides, and FFPE tissue sections were deparaffinized prior to Hematoxylin and eosin (H&E) or mucicarmine staining. Immunohistochemistry was performed on the Discovery Ultra (Ventana Medical Systems) using manufacturers reagents on 4μm sections. For anti-IGF-1 immunohistochemistry, anti-rabbit IGF-1 (Abcam #ab9572) was prepared using Discovery PSS Diluent (cat. #: 760-212). Antigen retrieval was performed using Ventana’s CC1 (pH 8.5) for 64 min at 90°C. The slides were given a hydrogen peroxide block for 8 min at room temperature and then incubated in the primary antibody diluent (1:100) for 1 hr at room temperature, followed by anti-Rat HRP secondary antibody (Ventana Omap OmniMap, #760-4457) for 32 min at room temperature. The slides were then treated with DAB and counterstained with Hematoxylin II for 12 min and then Bluing Reagent for 4 min.

#### Plasmids

The PGC-1α promoter constructs were generated by cloning the −500 to +52 bp of either the proximal or distal human PGC-1α promoter sequences via 5’ *NheI* and 3’ *HindIII* into the pGL4.15 luciferase reporter (Promega, #E6701). The human IGF-1 promoter constructs were generated by cloning either a 2079 bp region encompassing the P1 promoter or a 1547 bp region encompassing the P2 promoter via 5’ *BglII* and 3’ *HindIII* into either the pGL4.15 (Promega, #E6701) or the pGL3-Enhancer luciferase reporter (Promega, #E1771). The A-CREB (Ahn et al., 1998) dominant negative CREB cDNA was directionally cloned via 5′ *Not*I and 3′ *Mlu*I restriction enzyme sites into pRetroX-Tight-tdTomato (Amelio et al., 2014). The pTRE3G-MCS_PGK-GpNLuc construct was generated by excising the pTight promoter from the pRetroX-Tight-MCS_PGK-GpNLuc construct (Addgene plasmid #70185) containing our previously described LumiFluor optical reporter (Schaub et al., 2015) and subcloning the TRE3G promoter in its place. Subsequently, a N-terminal FLAG-tagged C1/M2 cDNA was directionally subcloned via 5’ *NotI* and 3’ *MluI* restriction enzyme sites into this pTRE3G-MCS-GpNLuc vector. The pLV-TRE3G_PGC1α4-mCherry:T2A:Hygro construct was custom synthesized (Cyagen Bioscience) to contain the human PPARGC1A variant 4 splice isoform (PGC-1α4) (Ruas et al., 2012) downstream of the TetO response element promoter for tet/dox inducible expression. The pFLAG-CMV-2_CRTC1/MAML2 (Tonon et al., 2003) construct was a gift from Frederic Kaye, the pLKO.1 shNS and shMAML2 (Chen et al., 2014) shRNA constructs were gifts from Lizi Wu, the pLMN UltramiR-E shIGF1R (ULTRA-3308043 and ULTRA-3443222) and shPPARGC1A (ULTRA-3197172 and ULTRA-3197171) constructs were from transOMICS Technologies, and the PPRE x3-TK-Luc (Kim et al., 1998) construct was a gift from Bruce Spiegelman.

#### PCR and qPCR

For cell lines and fresh-frozen tissues, gene expression was measured by extracting RNA using a Nucleospin RNA kit (Machery-Nagel #740955) according to the manufacturer’s instructions. cDNA was synthesized from 1 μg of RNA using iScript cDNA synthesis kit (Bio-Rad #170-8890). For human tissues, RNA was extracted using the Maxwell 16 MDx Instrument (Promega #AS3000) and the Maxwell 16 LEV RNA FFPE Kit (Promega #AS1260) according to the manufacturer’s protocol (Promega #9FB167). cDNA was made from 1-4 μg of RNA using SuperScript IV Reverse Transcriptase (Invitrogen #18090050) with dNTPs (NEB #N0446S), RNase inhibitor (Applied Biosystems #N808-0119), 25 μM oligo d(T)20 primer (Invitrogen #100023441) and 25 μM MAML2-specific reverse primer (Table S6). C1/M2 copy number was determined by establishing standard curves with 100 to 1×10^6^ copies of FLAG-C1/M2 plasmid. Relative gene expression of target genes was determined using the 2^ΔΔCt^ method and normalized to human *RPL23* expression. qPCR was performed using FastStart Universal SYBR Green Master (Rox) Mix (Roche #04913850001) with 1/50 (tissue) or 1/100 (cells) volume of the cDNA iScript reaction, and 0.25 μM of primers (Table S6).

#### Quantification and Statistical Analysis

All statistical tests were executed using GraphPad Prism software or the statistical software R (version 3.1.2). Differences between variables were assessed by 2-tailed Student’s t test or 2-way ANOVA with Bonferroni’s post hoc tests, where appropriate. Data are expressed as mean ± SEM, P values less than 0.05 were considered statistically significant (∗p < 0.05, ∗∗p < 0.01, ∗∗∗p < 0.001, ∗∗∗∗p < 0.0001).

#### SR10221 and SR2595 synthesis and purification

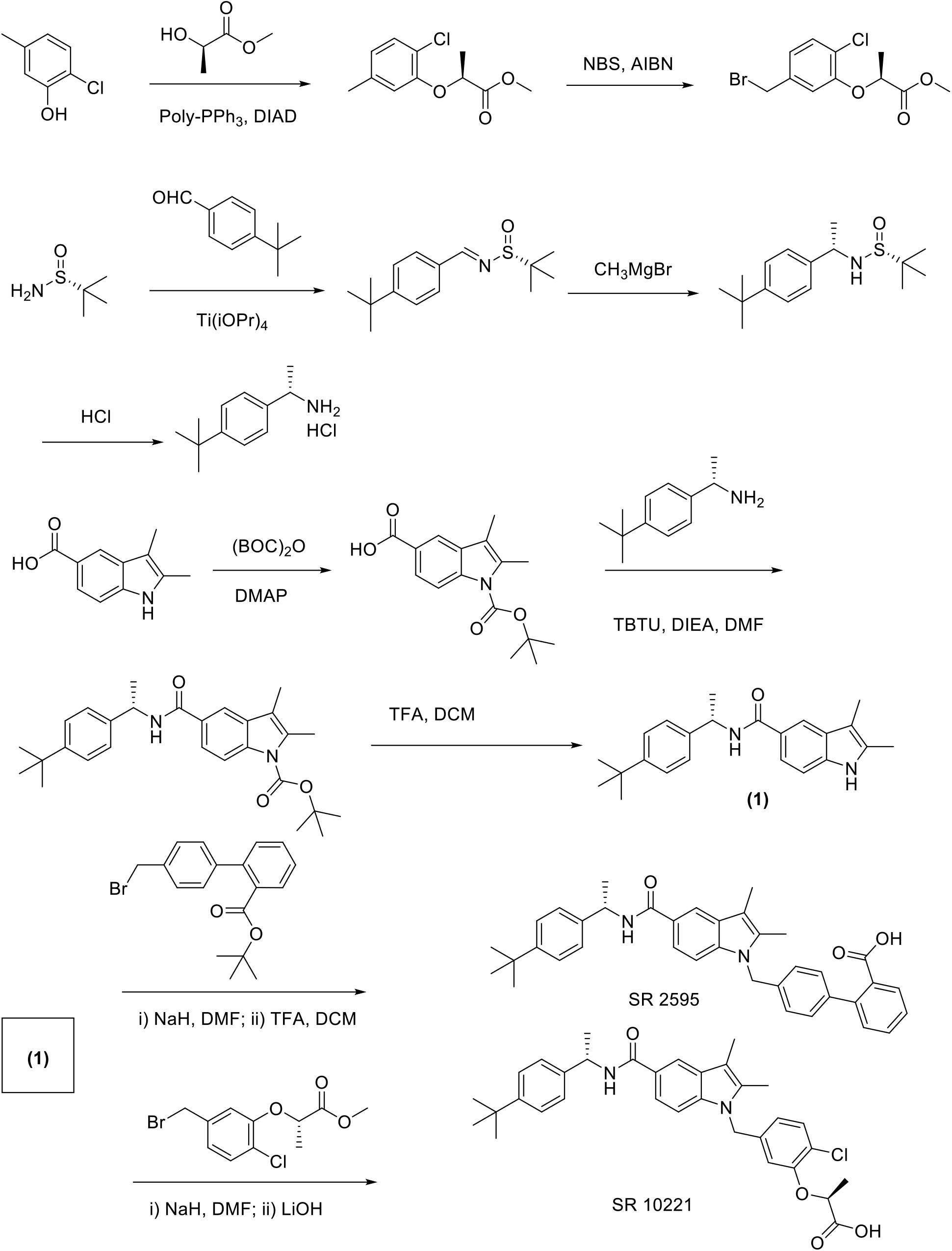

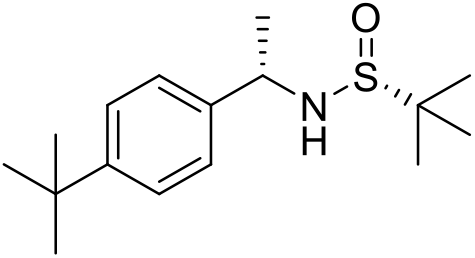

(R)-*N*-((S)-1-(4-(*tert*-butyl)phenyl)ethyl)-2-methylpropane-2-sulfinamide:

To a solution of 4-*t*-butylbenzaldehyde (1.03mL, 6.16mmol) in THF (50mL) (R)-2-methylpropane-2-sulfinamide (0.747g, 6.16mmol) was added. Titanium (IV) isopropoxide (9mL, 30.8mmol) was added dropwise to this solution and stirred for 20 h. It was then quenched with saturated solution of ammonium chloride and diluted with ethyl acetate. The mixture was filtered through a bed of celite and washed with ethyl acetate. The filtrate was separated in layers and the organic phases were combined and extracted with saturated solution of brine. It was then dried over anhydrous sodium sulfate, filtered and evaporated to dryness to obtain the imine as light yellow oil (1.54g). The imine showed 95% purity and used as is without any purification for the next step of reaction. Methylmagnesium bromide (3.9mL, 3.0M in diethyl ether) was added drop-wise to a solution of the crude imine (1.54g, 5.8mmol) in THF (50mL) at −50°C and the reaction was allowed to warm up to the room temperature for 18 h. It was then quenched with saturated brine and diluted with ethyl acetate. The organic phases were separated and dried over anhydrous sodium sulfate, filtered and evaporated to dryness. The crude residue was purified by chromatography on silica gel (Ethyl acetate/hexane) to obtain the title compound in 61% yield (1g).

1H NMR (400 MHz, DMSO-*d*6) δ ppm 1.11 (s, 9 H) 1.26 (s, 9 H) 1.43 (d, *J*=6.60 Hz, 3 H) 4.30 - 4.40 (m, 1 H) 5.26 (d, *J*=5.14 Hz, 1 H) 7.21 - 7.27 (m, 2 H) 7.29 - 7.38 (m, 2 H). ESI-MS (m/z) 282.1 [M+H]^+^.

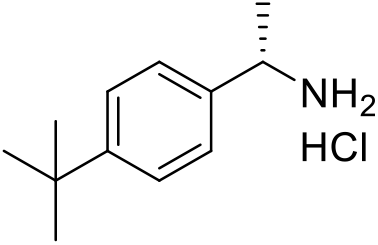

(S)-1-(4-(tert-butyl)phenyl)ethanaminiumchloride:

To a solution of (R)-*N*-((S)-1-(4-(*tert*-butyl)phenyl)ethyl)-2-methylpropane-2-sulfinamide 1g, 3.55mmol) in methanol (2mL) HCl solution (2mL, 4M in dioxane) was added and the reaction was monitored by looking at the disappearance of the starting material using LC-MS. When the strating material was all consumed, the reaction mixture was concentrated in vacuo. The crude residue was triturated in diethyl ether to get the title compound in 92% yield (0.7g).

1H NMR (400 MHz, DMSO-*d*6) δ ppm 1.28 (d, *J*=0.73 Hz, 9 H) 1.48 (d, *J*=6.85 Hz, 3 H) 4.34 (q, *J*=6.44 Hz, 1 H) 7.34 - 7.50 (m, 4 H) 8.25 (br. s, 3 H). ESI-MS (m/z) 161.1 [M+H-NH3]^+^.

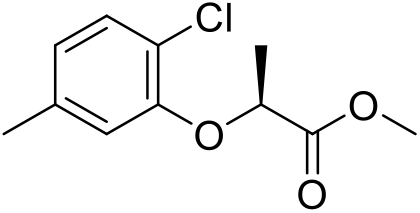

Methyl (S)-2-(2-chloro-5-methylphenoxy)propanoate:

To a solution of 2-chloro-5-methylphenol (0.5g, 3,5mmol) in anhydrous THF (20mL) at 0°C was added triphenylphosphine (1.38g, 5.25mmol) and methyl (R)-2-hydroxypropanoate 0.365g, 3.7mmol). Then diisopropyl azodicarboxylate (1.06g, 5.25mmol) was added drop-wise to the reaction at 0°C and stirred for 20 h. The solvent was removed and the crude was purified by chromatography on silica gel (Ethyl acetate/hexane) to obtain the title compound in 65% yield (0.525g).

1H NMR (400 MHz, CHLOROFORM-*d*) δ ppm 1.68 (s, 3 H) 2.30 (s, 3 H) 3.78 (s, 3 H) 4.76 (d, *J*=6.85 Hz, 1 H) 6.68 (s, 1 H) 6.71 - 6.80 (m, 1 H) 7.24 (d, *J*=8.07 Hz, 1 H). ESI-MS (m/z) 229.1 [M+H]^+^.

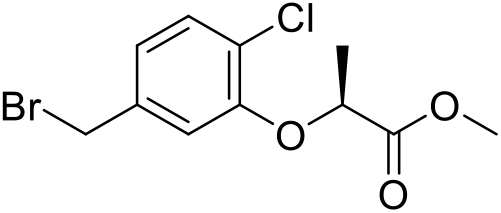

Methyl (S)-2-(5-(bromomethyl)-2-chlorophenoxy)propanoate:

To methyl (S)-2-(2-chloro-5-methylphenoxy)propanoate (0.55g, 2.4mmol) in dichloroethane (20mL) was added *N*-bromosuccinimide (0.428g, 2.4mmol) and azobisisobutyronitrile (0.039g, 0.24mmol). The mixture was refluxed for 2 h, cooled to room temperature and was concentrated in vacuo. The crude residue was purified by chromatography on silica gel (Ethyl acetate/hexane) to obtain the title compound in 78% yield (0.545g).

1H NMR (400 MHz, CHLOROFORM-*d*) δ ppm 1.68 - 1.72 (m, 3 H) 3.79 (d, *J*=0.98 Hz, 3 H) 4.42 (d, *J*=1.71 Hz, 2 H) 4.80 (d, *J*=6.85 Hz, 1 H) 6.88 (d, *J*=1.71 Hz, 1 H) 6.94 - 7.00 (m, 1 H) 7.34 (dd, *J*=8.07, 0.98 Hz, 1 H). ESI-MS (m/z) 306.1, 308.1 [M+H]^+^.

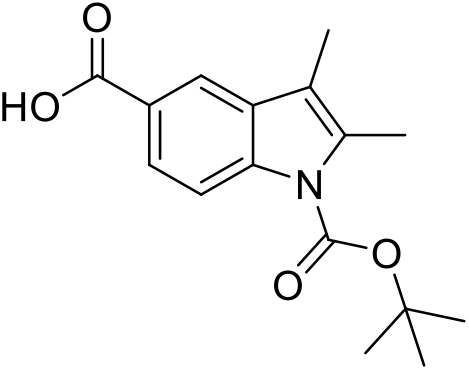

1-(tert-butoxycarbonyl)-2,3-dimethyl-1H-indole-5-carboxylic acid:

To a solution of 2,3-dimethyl-1H-indole-5-carboxylic acid (1.5g, 7.9mmol) in THF (50mL) was added di-*tert*-butyl dicarbonate (2.1g, 9.5mmol) and 4-(dimethylamino)pyridine (0.096g, 0.79mmol). The mixture was stirred for 20 h. It was then diluted with saturated ammonium chloride and ethyl acetate. The combined organic layers were separated and extracted with saturated brine, dried over anhydrous sodium sulfate, filtered and concentrated in vacuo. The crude residue was purified by chromatography on silica gel (Ethyl acetate/hexane) to obtain the title compound in 79% yield (1.8g).

1H NMR (400 MHz, DMSO-*d*6) δ ppm 1.65 (s, 9 H) 2.24 (s, 3 H) 2.52 (s, 3 H) 8.03 (d, *J*=8.80 Hz, 1 H) 8.21 (d, *J*=9.05 Hz, 1 H) 8.29 (s, 1 H). ESI-MS (m/z) 190.1 [M+H-100]^+^.

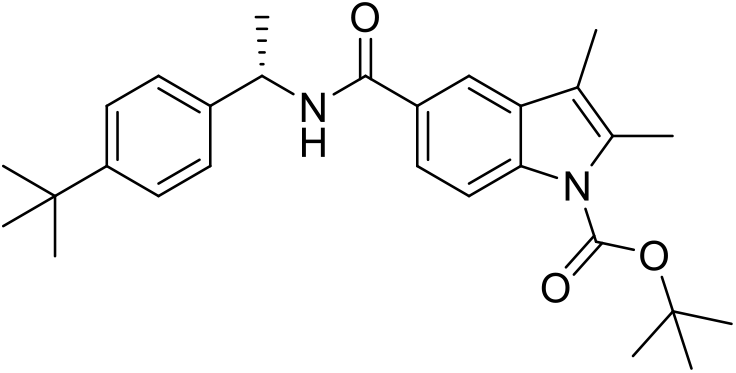

*tert-*Butyl-(S)-5-((1-(4-(tert-butyl)phenyl)ethyl)carbamoyl)-2,3-dimethyl-1H-indole-1-carboxylate: To a mixture of 1-(tert-butoxycarbonyl)-2,3-dimethyl-1H-indole-5-carboxylic acid (0.2g, 0.69mmol) and (S)-1-(4-(tert-butyl)phenyl)ethanaminiumchloride (0.178g, 0.83mmol) in dimethylformamide (5mL) was added *N,N*-diisopropylethylamine (360μL, 2.07mmol) and *O*-(benzotriazol-1yl)-*N,N,N’,N’*-tetramethyluronium tetrafluroborate (0.222g, 0.69mmol) and the mixture was stirred for 20 h. It was then diluted with saturated solution of sodium bicarbonate and ethyl acetate. The combined organic layers were separated and extracted with saturated brine, dried over anhydrous sodium sulfate, filtered and concentrated in vacuo. The crude residue was purified by chromatography on silica gel (Ethyl acetate/hexane) to obtain the title compound in 87% yield (0.269g).

1H NMR (400 MHz, DMSO-*d*6) δ ppm 1.28 (s, 9 H) 1.49 (d, *J*=7.09 Hz, 3 H) 1.63 (s, 9 H) 2.21 (d, *J*=0.73 Hz, 3 H) 5.08 - 5.25 (m, 1 H) 7.28 - 7.39 (m, 4 H) 7.80 (dd, *J*=8.68, 1.83 Hz, 1 H) 7.96 – 8.09 (m, 2 H) 8.73 (d, *J*=8.07 Hz, 1 H). ESI-MS (m/z) 349.1 [M+H-100]^+^.

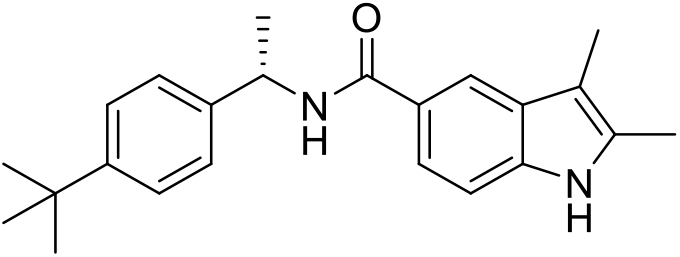

(S)-*N*-(1-(4-(tert-butyl)phenyl)ethyl)-2,3-dimethyl-1H-indole-5-carboxamide:

To a mixture of *tert-*Butyl-(S)-5-((1-(4-(tert-butyl)phenyl)ethyl)carbamoyl)-2,3-dimethyl-1H-indole-1-carboxylate (0.24g, 0.53mmol) in dichloromethane (5mL) was added trifluroacetic acid (0.5mL) at 0°C and the mixture was stirred for 0.5 h. It was then concentrated in vacuo, diluted with saturated solution of sodium bicarbonate and ethyl acetate. The combined organic layers were separated and extracted with saturated brine, dried over anhydrous sodium sulfate, filtered and concentrated in vacuo. The crude residue was triturated in diethyl ether to get the title compound in 99% yield (0.7g) in 90% purity. The free base was used as is without any purification for the next step of reaction. ESI-MS (m/z) 349.1 [M+H]^+^.

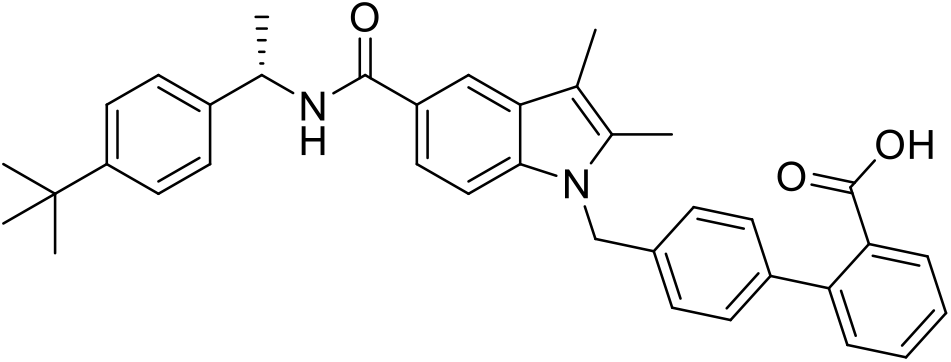

##### SR2595

(S)-4’-((5-((1-(4-(tert-butyl)phenyl)ethyl)carbamoyl)-2,3-dimethyl-1H-indol-1-yl)methyl)-[1,1’-biphenyl]-2-carboxylic acid:

To a solution of (S)-*N*-(1-(4-(tert-butyl)phenyl)ethyl)-2,3-dimethyl-1H-indole-5-carboxamide (0.05g, 0.14mmol) and *tert*-butyl 4’-(bromomethyl)-[1,1’-biphenyl]-2-carboxylate (0.059g, 0.17mmol) in anhydrous dimethylformamide (2mL) at 0°C was added sodium hydride (0.007g, 0.228mmol) and the mixture was stirred for 0.5 h. It was then quenched with methanol and concentrated in vacuo. The crude was then taken in dichloro methane (2mL) and trifluroacetic acid (0.25mL) was added and stirred for 0.5 h. The mixture was concentrated in vacuo and the crude residue was purified by reversed phase chromatography (water/acetonitrile) to obtain the title compound in 41% yield (0.032g).

1H NMR (400 MHz, DMSO-*d*6) δ ppm 8.59 (d, *J =* 8.0 Hz, 1 H), 8.10 (d, *J =* 1.5 Hz, 1 H), 8.68 (dd, *J =* 1.0, 7.6 Hz, 1 H), 7.63 (dd, *J =* 1.5, 8.5 Hz, 1 H), 7.51 (m, 1 H), 7.44 (d, *J =* 8.8 Hz, 1 H), 7.40 (dd, *J =* 1.2, 7.6 Hz, 1 H), 7.31-7.33 (m, 4 H), 7.23 (d, *J =* 8.3 Hz, 2 H), 6.98 (d, *J =* 8.3 Hz, 2 H), 5.47 (s, 2H), 5.33 (m, 1 H), 2.32 (s, 3 H), 2.28 (s, 3 H), 1.26 (s, 9 H). ESI-MS (m/z) 559.1 [M + H]^+^.

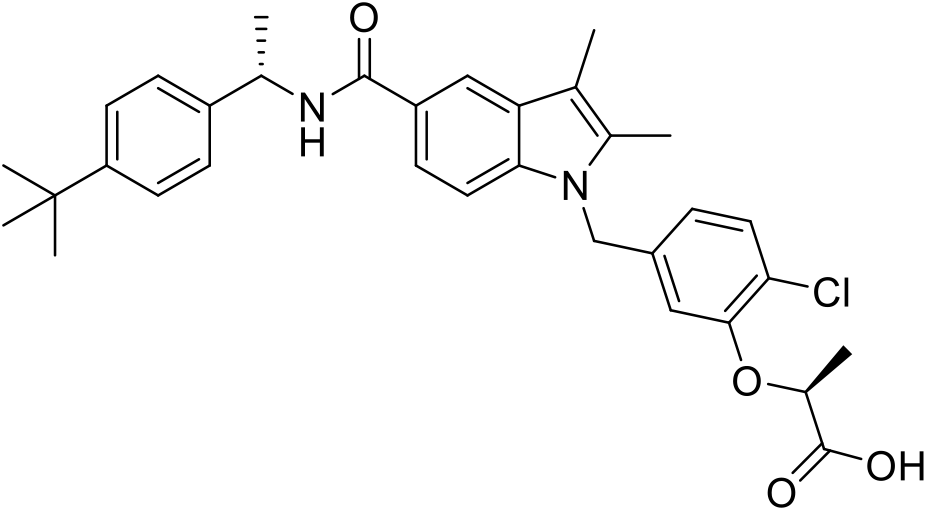

##### SR10221

(S)-2-(5-((5-(((S)-1-(4-(tert-butyl)phenyl)ethyl)carbamoyl)-2,3-dimethyl-1H-indol-1-yl)methyl)-2-chlorophenoxy)propanoic acid:

To a solution of (S)-*N*-(1-(4-(tert-butyl)phenyl)ethyl)-2,3-dimethyl-1H-indole-5-carboxamide (0.05g, 0.14mmol) and methyl (S)-2-(5-(bromomethyl)-2-chlorophenoxy)propanoate (0.053g, 0.17mmol) in anhydrous dimethylformamide (2mL) at 0°C was added sodium hydride (0.007g, 0.228mmol) and the mixture was stirred for 0.5 h. It was then quenched with methanol and concentrated in vacuo. The crude was then taken in aqueous lithium hydroxide solution (2mL) and methanol (0.25mL) was added and stirred for 0.5 h. The mixture was concentrated in vacuo and the crude residue was purified by reversed phase chromatography (water/acetonitrile) to obtain the title compound in 38% yield (0.030g).

1H NMR (400 MHz, DMSO-*d*6) δ ppm 8.57 (d, *J*=8.1 Hz, 1 H), 8.08 (d, *J*=1.5Hz, 1 H), 7.60 (dd, *J*=1.8, 8.6 Hz, 1 H), 7.36 (d, *J*=8.6 Hz, 1 H), 7.33 (m, 3 H), 7.29 (d, *J*=.1 Hz, 1 H), 6.79 (d, *J*=1.8 Hz, 1 H), 6.34 (dd, *J*=1.8, 8.1Hz, 1 H), 5.47 (s, 2H), 5.17 (m, 1 H), 4.78 (m, 1H), 2.27 (s, 3 H), 2.26 (s, 3 H), 1.50(d, *J*=6.8 Hz, 3 H), 1.47 (d, *J*=7.1 Hz, 3H), 1.26 (s, 9 H).). ESI-MS (m/z) 562.1 [M + H]^+^.

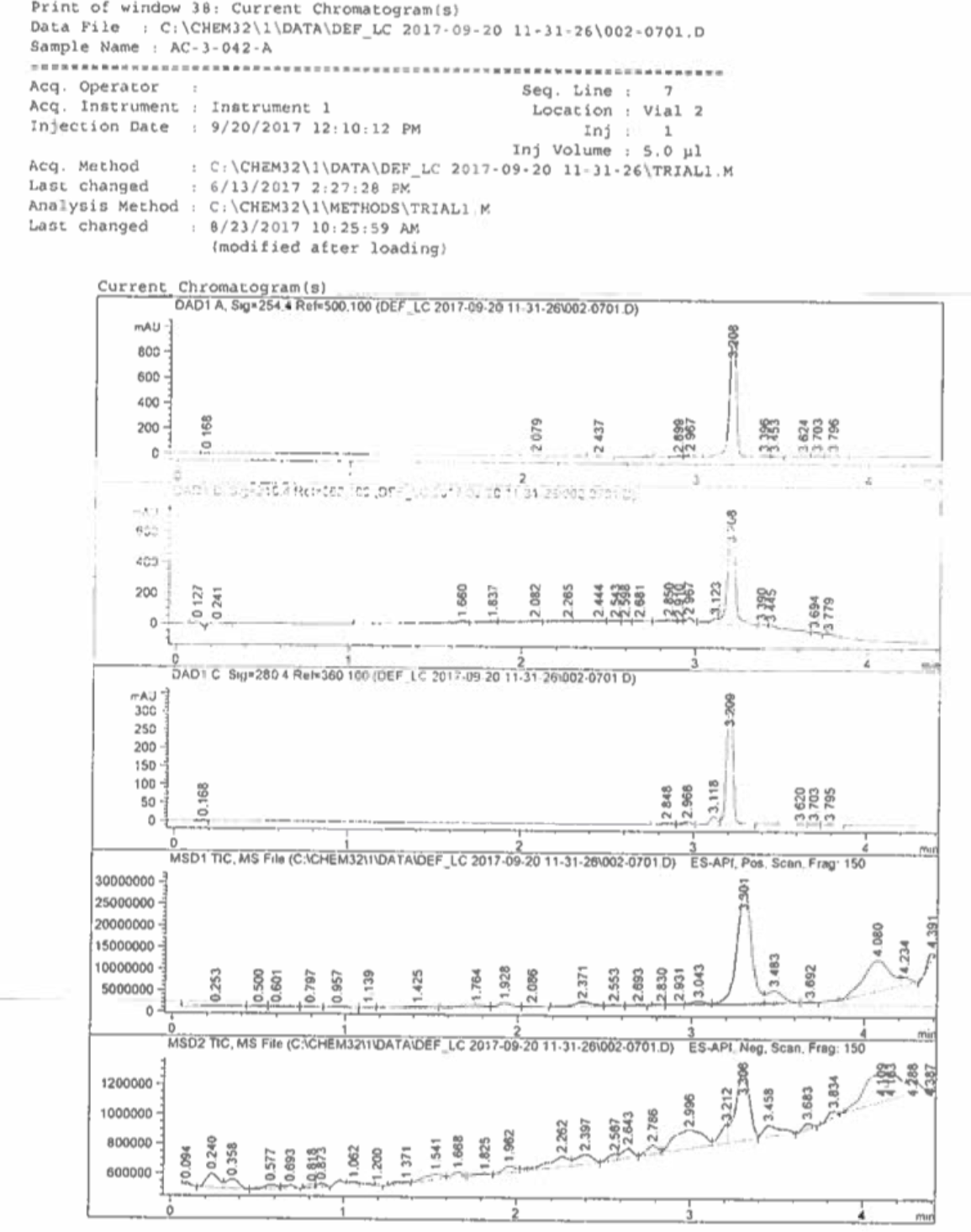
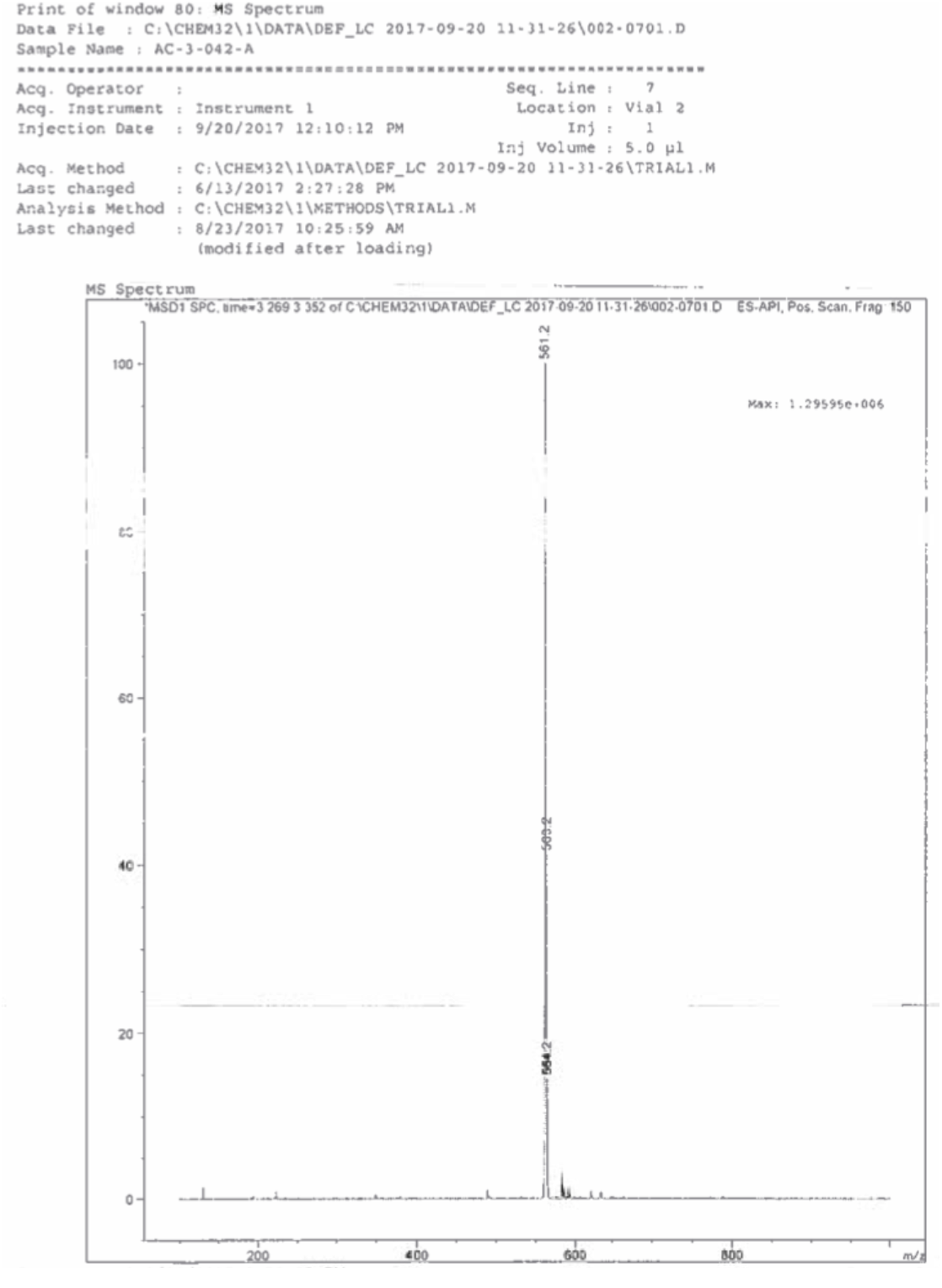

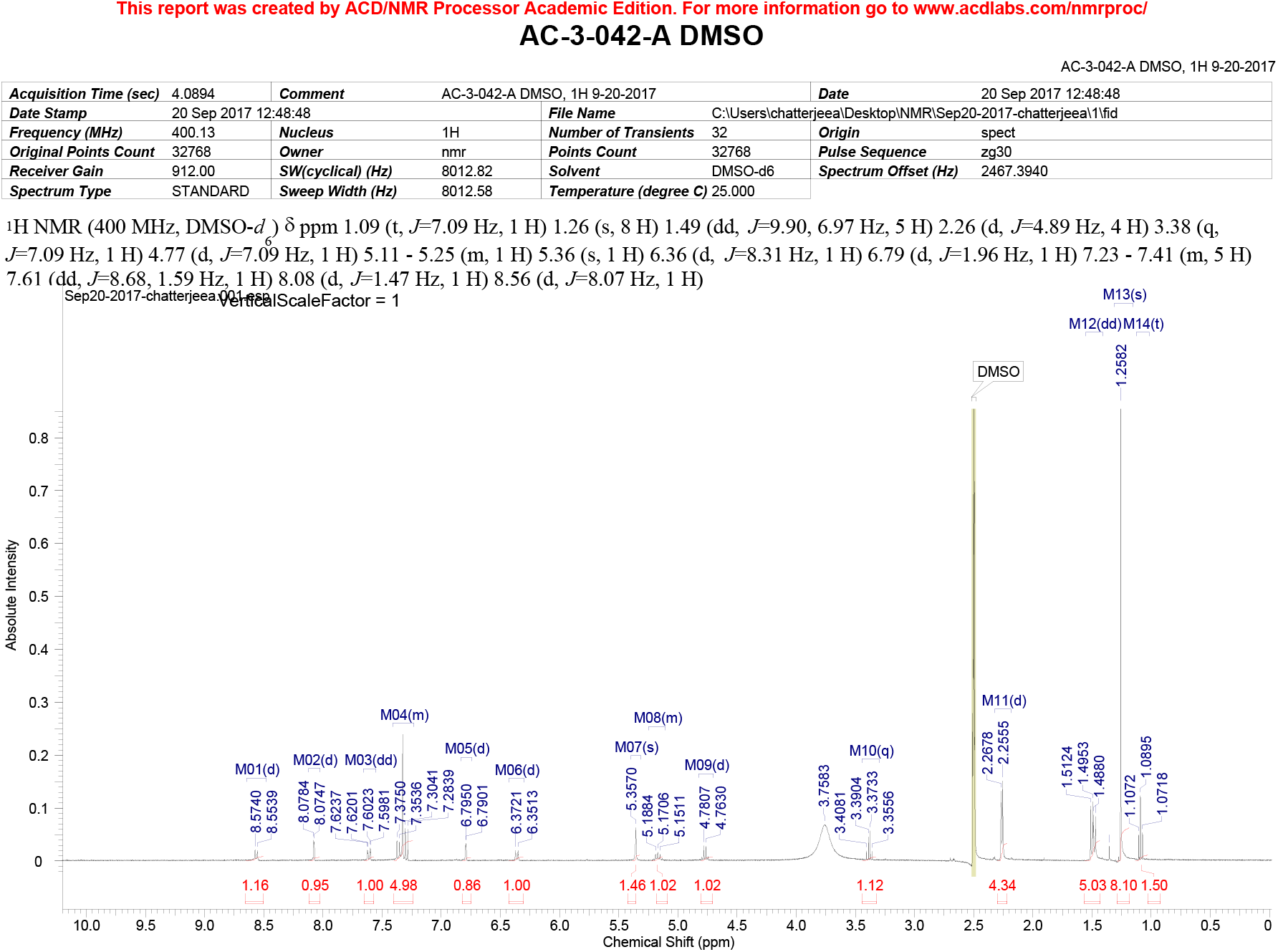

---

**Table S1. Cancer Pathway Genes induced upon CRTC1-MAML2 expression. Related to Figures 1 and S1.**

**Table S2A. Differentially expressed genes in salivary MEC tumors versus normal salivary gland tissues. Related to Figure S1.**

**Table S2B. Top 1500 differentially expressed genes in salivary MEC tumors versus normal salivary gland tissues. Related to Figure S1.**

**Table S3. Differentially expressed genes in C1/M2-positive salivary MEC tumors versus normal salivary gland tissues. Related to Figure 1.**

**Table S4. Curated list of differentially expressed IGF-PI3K pathway genes induced by C1/M2. Related to Figures 1 and S1.**

**Table S5. Genomics of IGF1R-PI3K inhibitor Drug Sensitivity in Salivary Cancer Cells. Related to Figures 2 and S3.**

**Table S6. List of Primers. Related to STAR Methods.**

