## Supplementary Information for "CRTC1-MAML2 Establishes a PGC1α-IGF1 Circuit that Confers Vulnerability to PPARγ Inhibition"

[1]Graduate Curriculum in Genetics and Molecular Biology, Biological and Biomedical Sciences Program, UNC School of Medicine, University of North Carolina at Chapel Hill, Chapel Hill, NC, USA
[2]Graduate Curriculum in Cell Biology and Physiology, Biological and Biomedical Sciences Program, UNC School of Medicine, University of North Carolina at Chapel Hill, Chapel Hill, NC, USA
[3]Bioinformatics Core, Lineberger Comprehensive Cancer Center, UNC School of Medicine, University of North Carolina at Chapel Hill, Chapel Hill, NC, USA
[4]Graduate Curriculum in Pharmacological and Physiological Sciences, School of Medicine, Saint Louis University, Saint Louis, MO, USA
[5]Department of Pharmacology and Physiology, School of Medicine, Saint Louis University, Saint Louis, MO, USA
[6]Division of Oral and Craniofacial Health Sciences, UNC Adams School of Dentistry, University of North Carolina at Chapel Hill, Chapel Hill, NC, USA
[7]Lineberger Comprehensive Cancer Center, Cancer Genetics Program, UNC School of Medicine, University of North Carolina at Chapel Hill, Chapel Hill, NC, USA
[8]Division of Hematology/Oncology, Department of Medicine, UNC School of Medicine, University of North Carolina at Chapel Hill, Chapel Hill, NC, USA
[9]Department of Pathology and Laboratory Medicine, UNC School of Medicine, University of North Carolina at Chapel Hill, Chapel Hill, NC, USA
[10]Department of Otolaryngology/Head and Neck Surgery, UNC School of Medicine, University of North Carolina at Chapel Hill, Chapel Hill, NC, USA
[11]Division of Diagnostic Sciences, UNC Adams School of Dentistry, University of North Carolina at Chapel Hill, Chapel Hill, NC, USA
[12]Department of Genetics, UNC School of Medicine, University of North Carolina at Chapel Hill, Chapel Hill, NC, USA
[13]Department of Cell Biology and Physiology, UNC School of Medicine, University of North Carolina at Chapel Hill, Chapel Hill, NC, USA
[14]McAllister Heart Institute, UNC School of Medicine, University of North Carolina at Chapel Hill, Chapel Hill, NC, USA
[15]Department of Medical Oncology, University of Tennessee Health Sciences West Cancer Center, Memphis, Tennessee, USA
[16]Biomedical Research Imaging Center, UNC School of Medicine, University of North Carolina at Chapel Hill, Chapel Hill, NC, USA
[17]Lineberger Comprehensive Cancer Center, Cancer Cell Biology Program, UNC School of Medicine, University of North Carolina at Chapel Hill, Chapel Hill, NC, USA
[18]Lead Contact

**Supplementary Information Includes**:

Supplemental Figures S1-S6
Supplemental Tables S1-S6

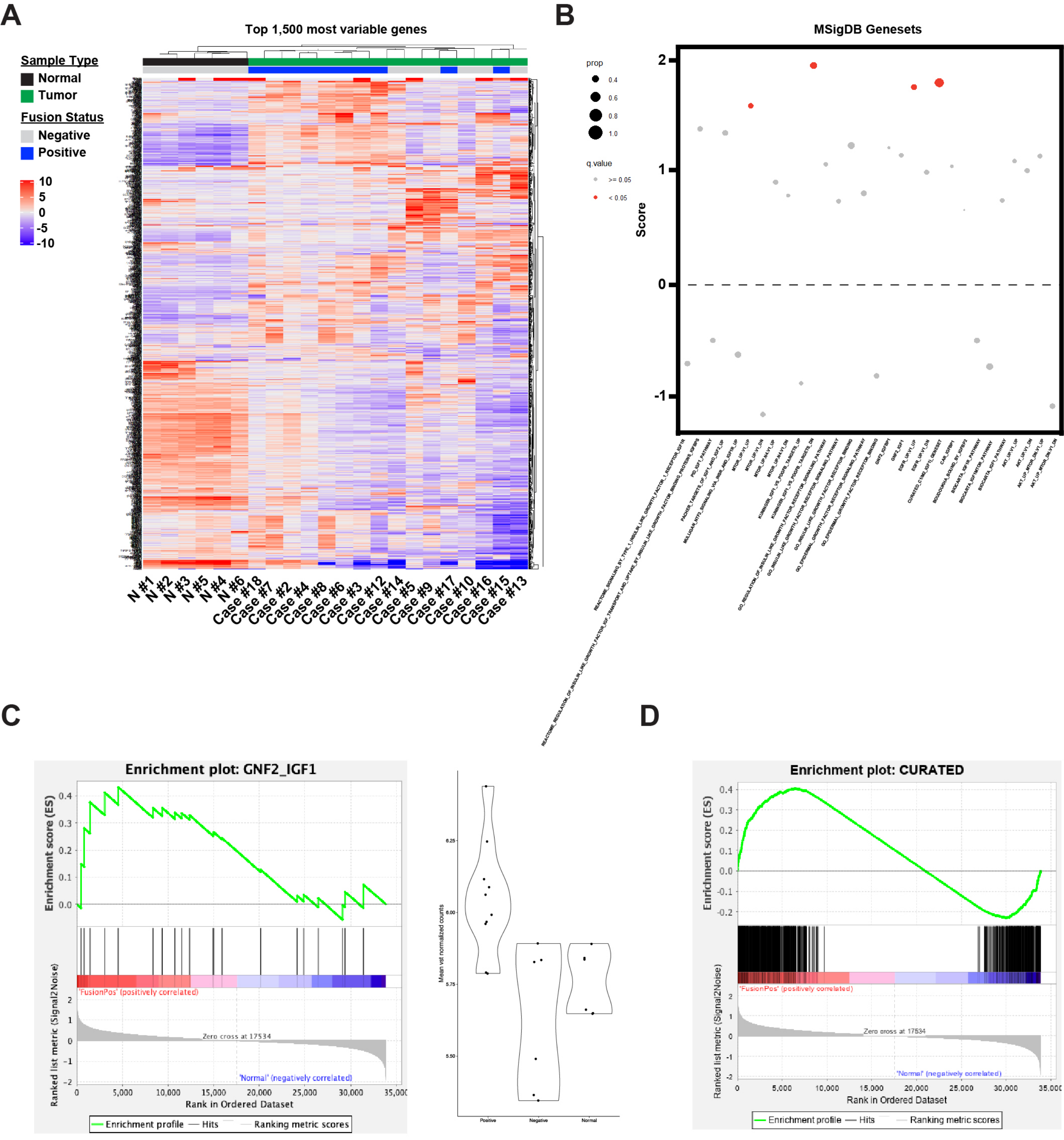


### Supplemental Figure S1

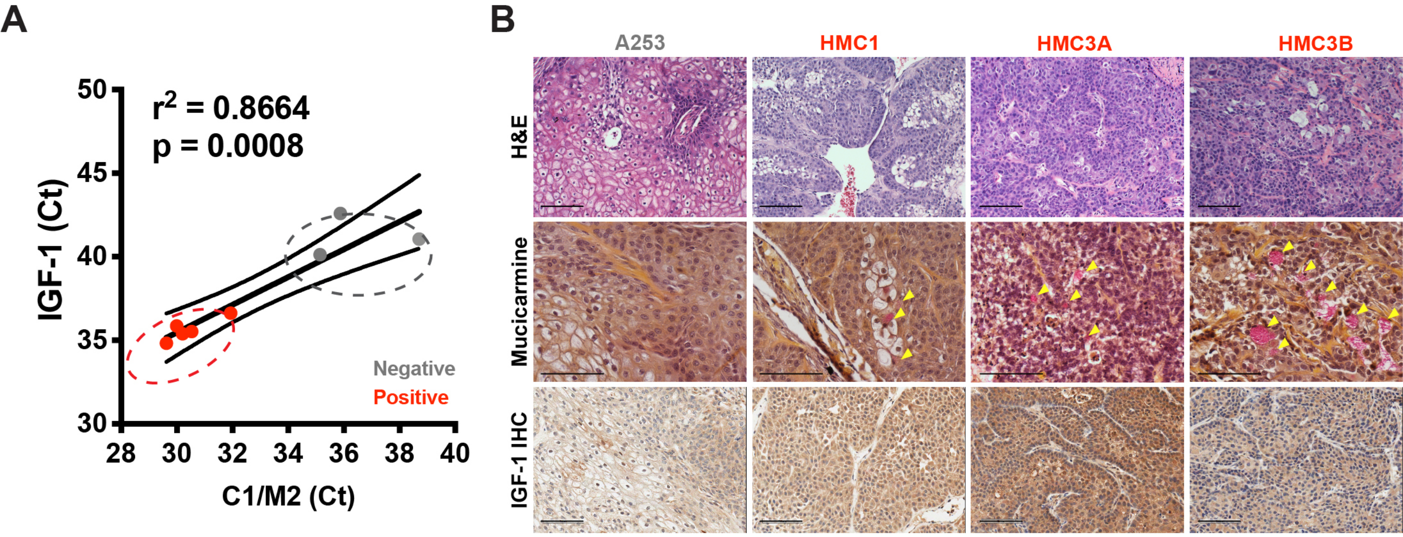


**Supplemental Figure S2**

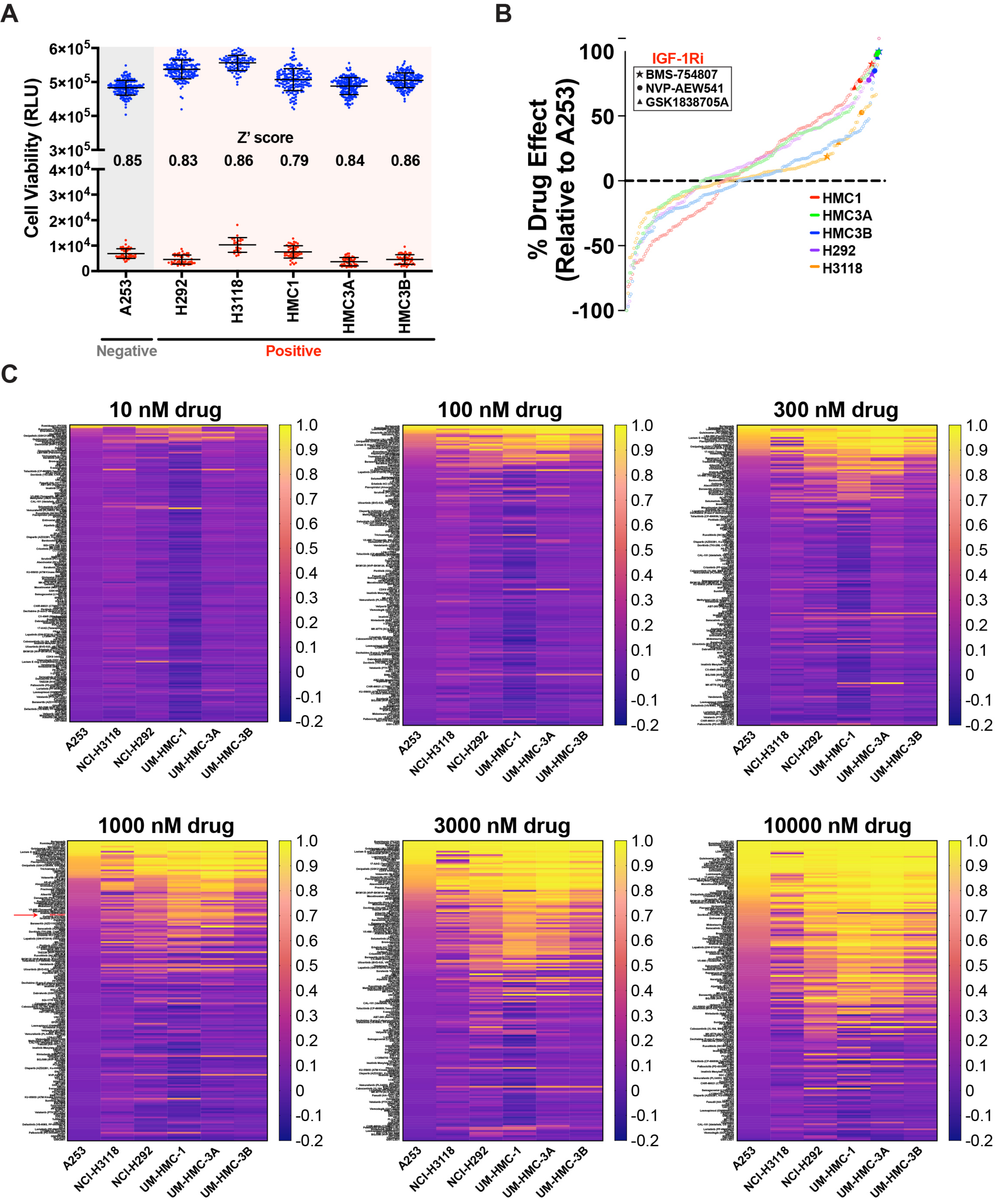


Supplemental Figure S3

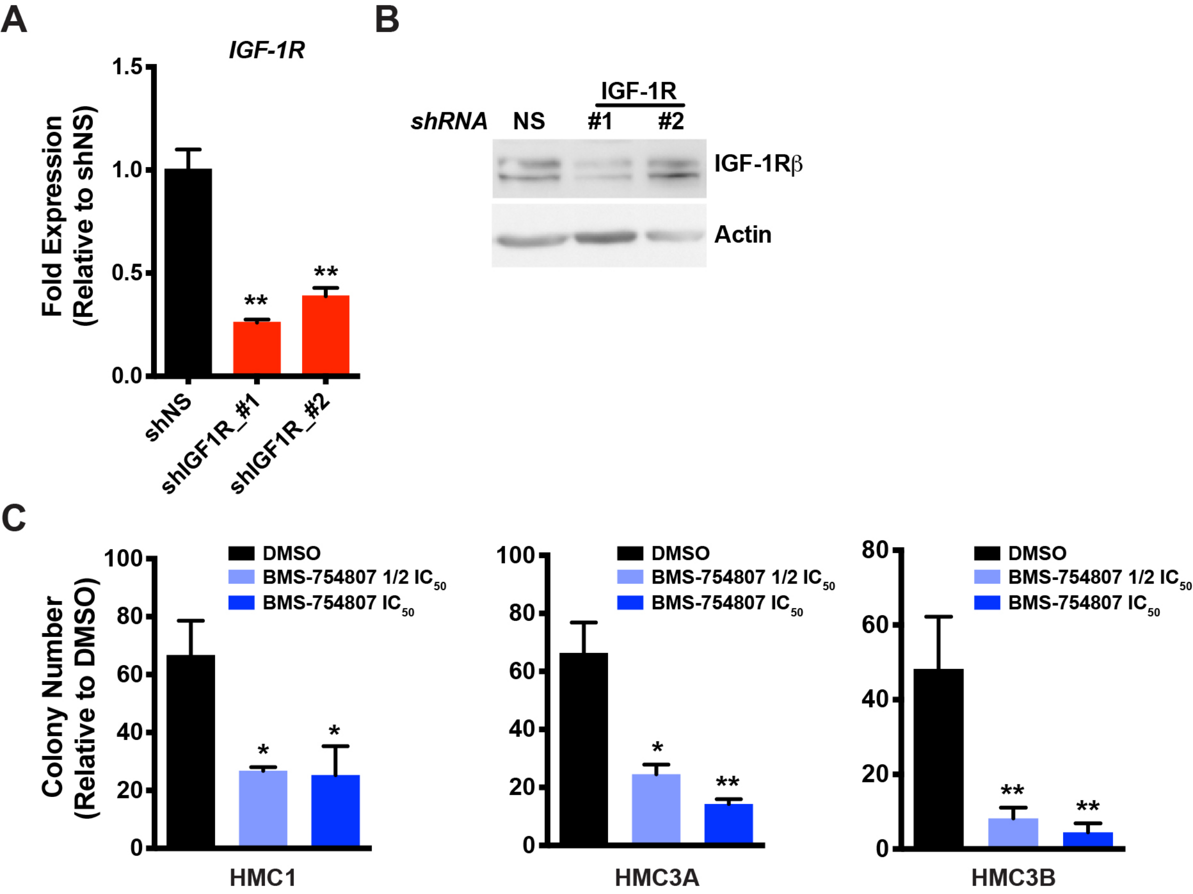


Supplemental Figure S4

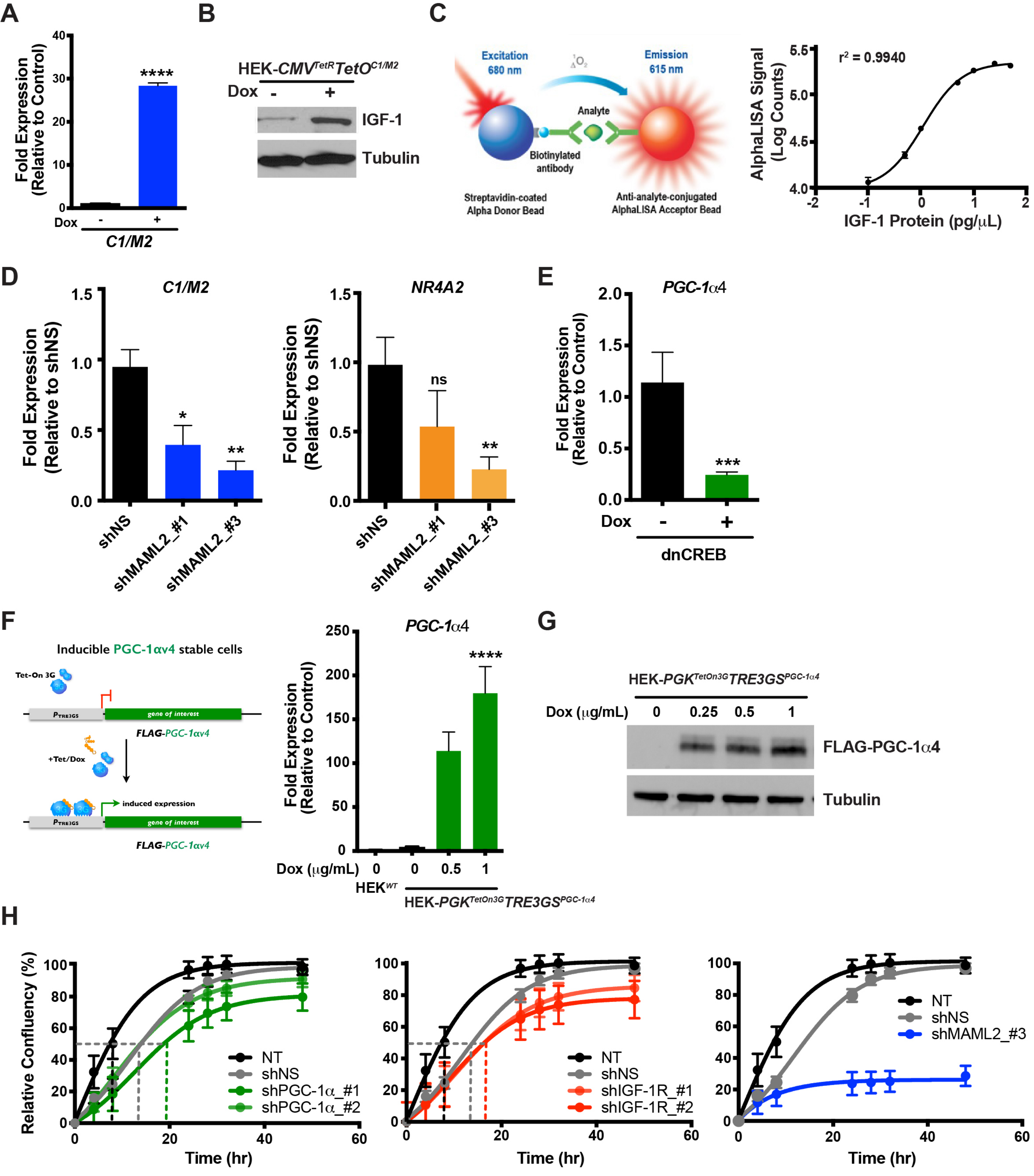


#### Supplemental Figure S5

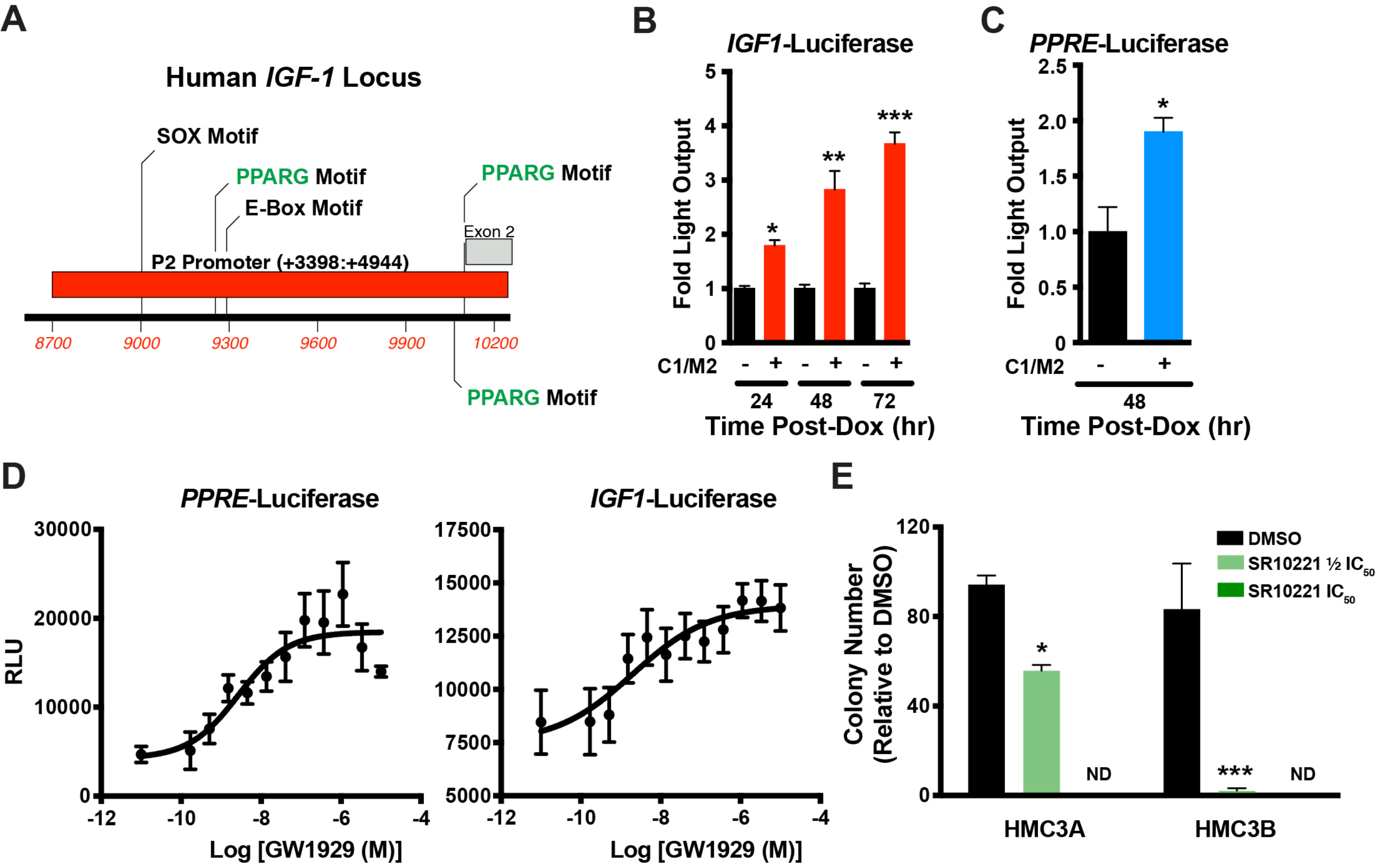


### Supplemental Figure S6
